# Visualizing conformational space of functional biomolecular complexes by deep manifold learning

**DOI:** 10.1101/2021.08.09.455739

**Authors:** Zhaolong Wu, Enbo Chen, Shuwen Zhang, Yinping Ma, Congcong Liu, Chang-Cheng Yin, Youdong Mao

## Abstract

The cellular functions are executed by biological macromolecular complexes in nonequilibrium dynamic processes, which exhibit a vast diversity of conformational states. Solving conformational continuum of important biomolecular complexes at atomic level is essential to understand their functional mechanisms and to guide structure-based drug discovery. Here we introduce a deep learning framework, named AlphaCryo4D, which enables atomic-level cryogenic electron microscopy reconstructions of conformational continuum. AlphaCryo4D integrates 3D deep residual learning with manifold embedding of energy landscapes, which simultaneously improves 3D classification accuracy and reconstruction resolution via an energy-based particle-voting algorithm. By applications of this approach to analyze five experimental datasets, we demonstrate its generality in breaking resolution barrier of visualizing dynamic components of functional complexes, in choreographing continuous inter-subunit motions and in exploring their ‘hidden’ conformational space. Our approach expands the realm of structural ensemble determination to the nonequilibrium regime at atomic level, thus potentially transforming biomedical research and therapeutic development.

---

Essential cellular functions are executed and regulated by macromolecular complexes comprising many subunits, such as the ribosome, proteasome, replisome and spliceosome. Their structures in functional cycles are often compositionally heterogeneous and conformationally dynamic, involving many reversibly associated components in cells. Visualizing the conformational continuum of highly dynamic megadalton complexes at high resolution is crucial for understanding their functional mechanisms and for guiding structure-based therapeutic development. The current approaches in cryogenic electron microscopy (cryo-EM) structure determination allow atomic-level visualization of major conformational states of these dynamic complexes^1–12^. However, high-resolution structure determination of transient, nonequilibrium intermediates connecting their major states during their functional cycles has been prohibitory at large. To date, there has been lack of appropriate approaches in visualizing conformational space of large biomolecular complexes at high resolution.

The problem of structural heterogeneity in cryo-EM structure determination has been investigated with numerous computational approaches including maximum-likelihood-based classification and multivariate statistical analysis^5, 6, 11, 13–18^. As an extension of multivariate statistical analysis, several machine-learning approaches were recently proposed to estimate a continuous conformational distribution in the latent space^11, 12, 19–22^. The estimation of continuous conformational distributions does not guarantee an improvement of 3D classification accuracy nor warrant identification of hidden conformational states of biological importance. These methods often trade off the reconstruction resolution for gaining an overall representation of conformational landscape in the latent space. The insufficient resolution would preclude their potential applications in structure-based drug discovery.

To date, the most widely used method to counteract structural heterogeneity in cryo-EM for resolution improvement is the hierarchical maximum-likelihood-based 3D classification due to its ease of application^1^. To improve the resolution of major conformers, low-quality classes are often manually removed during iterations of curated hierarchical classification, inevitably causing an incomplete representation of the true conformational landscape^23, 24^. The outcome of refined cryo-EM maps depends on user expertise of decision-making in class selection, which can be biased by user experience and subjectivity. Importantly, a considerable portion of misclassified images can limit the achievable resolution of observed conformational states and result in missing states that could be biologically important.

In addition to the approaches characterizing global structural heterogeneity, other methods were proposed to analyze local structural dynamics or to improve local resolution of flexible regions in cryo-EM reconstructions. These includes the multi-body or focused refinement^4, 25, 26^, normal mode analysis^27, 28^, flexible refinement^29, 30^, non-uniform refinement^31^ and integrating traditional molecular dynamics (MD) simulations with or without use of deep learning^32, 33^. These methods were not designed to recover a complete picture of structural dynamics hidden in cryo-EM datasets.

The energy landscape is a statistical physical representation of the conformational space of a macromolecular complex and is the basis of the transition-state theory of chemical reaction dynamics^34, 35^. The minimum-energy path (MEP) on the energy landscape theoretically represents the most probable trajectory of conformational transitions and can inform the activation energy for chemical reactions^35, 36^. Previous studies have demonstrated the benefit of energy landscape estimation in characterizing conformational variation of macromolecules from cryo-EM data^37–43^. The combination of linear dimensionality reduction via 3D principal component analysis (PCA) and energy landscape estimation has been demonstrated to be capable of capturing the overall conformational landscape^38–40^. The benefit of using nonlinear dimensionality reduction and manifold embedding, such as diffusion map^44^, to replace PCA for estimating energy landscapes has also been investigated at limited resolution, with the assumption that the conformational changes can be discerned through a narrow angular aperture^37, 41, 42^. This assumption restricts its potential applications to more complicated conformational dynamics.

Breaking the resolution barrier in visualizing the conformational space of highly dynamic complexes represent a major objective of technical advances for leveraging cryo-EM applications in both basic biomedical science and drug discovery. This would require substantial improvement in 3D classification techniques that are optimized for resolution improvement. We hypothesize that a high-quality estimation of energy landscape could be potentially used to improve 3D classification of cryo-EM data. If different conformers or continuous conformational changes can be sufficiently mapped and differentiated on the energy landscape, the energetic visualization of conformational continuum could then be used to discover previously ‘invisible’ conformers and achieve higher resolution for lowly populated, transient or nonequilibrium states. To explore these ideas, we developed a novel machine learning framework named AlphaCryo4D that can break the existing limitations and enable 4D cryo-EM reconstruction of highly dynamic, lowly populated intermediates or transient states at atomic level. We examined the general applicability of AlphaCryo4D in analyzing several large synthetic datasets over a wide range of single-to-noise ratios (SNRs) and five experimental cryo-EM datasets of diverse sample behaviors in complex dynamics, including the mammalian calcium-conducting channel RyR1 complex^45^, human 26S proteasome^23^, malaria parasite 80S ribosome^46^, pre-catalytic spliceosome^47^ and bacterial ribosomal assembly intermediates^48^. Our approach pushes the envelope of visualizing conformational space of macromolecular complexes beyond the previously achieved scope toward the atomic level.

## Results

### Design of deep manifold learning

The conceptual framework of AlphaCryo4D integrates unsupervised deep learning with manifold embedding to learn a free-energy landscape, which directs cryo-EM reconstructions of conformational continuum or transient states via an energy-based particle-voting algorithm. AlphaCryo4D consists of four major steps (Fig. 1a). First, hundreds to thousands of 3D volumes are bootstrapped with *M*-fold particle shuffling and Bayesian clustering. To this end, all curated particles are aligned in a common frame of reference through a consensus 3D refinement. They are then randomly divided into *M*+1 groups of equal data sizes. In the step of ‘*M*-fold particle shuffling’, one group of particles is taken out of the dataset to form a shuffled dataset. This procedure is repeated *M* + 1 times, each time with a different particle group being left out, resulting in *M* + 1 shuffled datasets (Supplementary Fig. 1). Each shuffled dataset is subject to 3D volume bootstrapping separately and is clustered into tens to hundreds of 3D reconstructions through Bayesian clustering in RELION ^3, 7^. In total, hundreds to thousands of volumes from all shuffled datasets are expected to be bootstrapped through these steps.

**Fig. 1.**
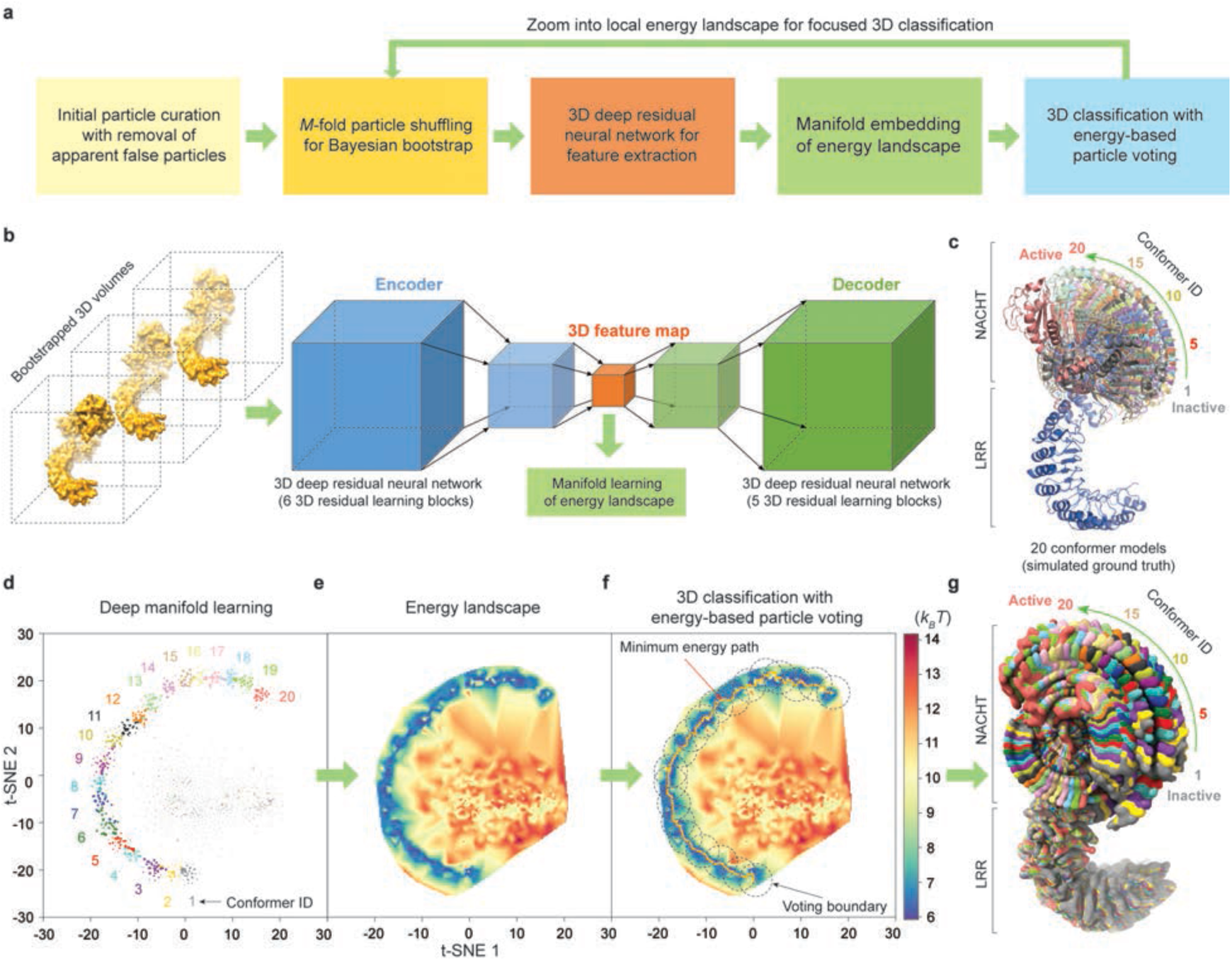
Conceptual framework of AlphaCryo4D for 4D cryo-EM reconstruction. **a,** Schematic showing the major conceptual steps of single-particle cryo-EM data processing in AlphaCryo4D. **b,** Illustration of deep residual learning of 3D feature maps by an autoencoder conjugated to a decoder for unsupervised training. **c,** The 20 atomic models of hypothetical conformers of NLRP3 in cartoon representations simulate a 90° rotation of the NACHT domain relative to the LRR domain based on the structure of NLRP3 (PDB ID: 6NPY). **d,** Manifold learning of the bootstrapped 3D volumes and their feature maps learnt by the autoencoder. Each data point corresponds to a 3D volume. The color labels the conformer identity of ground truth for the purpose of verification. **e,** Energy landscape computed from the manifold shown in (**d**) using the Boltzmann distribution. **f,** Minimum energy path (orange line) calculated by the string method is used to find the approximate cluster centers of 20 conformers. The cluster boundaries for energy-based particle voting are shown as dashed circles. **g,** The 20 reconstructions resulting from the blind assessment of AlphaCryo4D using simulated dataset with SNR of 0.01 are superimposed together, shown in surface representation, and aligned against its LRR domain.

Next, all bootstrapped volumes are learned by a 3D autoencoder made of a deep residual convolutional neural network in an unsupervised fashion^49, 50^ (Fig. 1b, Supplementary Table 1). A 3D feature map is extracted for each volume by the deep residual autoencoder and is juxtaposed with the volume data for nonlinear dimensionality reduction by manifold embedding with the t-distributed stochastic neighbor embedding (t-SNE) algorithm^51^ (Fig. 1d, e).

Third, the learned manifold is used to estimate an energy landscape via the Boltzmann relation using the particle density on the manifold^37^. A string method can be optionally used to search the MEP on the energy landscape^52, 53^. The local energy minima or transition states connecting adjacent minimum-energy states can be defined as the centers of 3D clustering, with a circular range defined as the cluster boundary for subsequent particle voting (Fig. 1f).

Last, because each particle is used *M* times during volume bootstrapping, it is mapped to *M* locations on the energy landscape. The mapping of each copy of the particle is called a ‘vote’. By counting the number of votes of the same particle casted within the same cluster boundary on the energy landscape, the reproducibility of machine learning procedure can be evaluated at single-particle level. Each particle is classified to the 3D cluster that receives more than *M*/2 votes of this particle within its voting boundary (Fig. 1f, Supplementary Fig. 1b). If none of the 3D clusters on the energy landscape receives more than *M*/2 votes of a given particle, the corresponding particle is voted out and excluded for further 3D reconstruction. The resulting 3D classes are expected to be conformationally homogeneous enough for high-resolution cryo-EM refinement.

Since many protein complexes exhibit profound conformational changes in different local regions, we also implemented a focused classification strategy of AlphaCryo4D that applies a local 3D mask^6^ throughout the entire procedure, which is executed as an iterative step after initial 3D classification by AlphaCryo4D in the absence of any 3D mask (Fig. 1a, Methods).

### Solving conformational continuum at atomic level

To assess the numerical performance of AlphaCryo4D, we generated three large synthetic heterogeneous cryo-EM datasets with signal-to-noise ratios (SNRs) of 0.05, 0.01 and 0.005. Each dataset includes 2 million randomly oriented single particles computationally simulated from 20 hypothetical conformer models of the ∼130-kDa NLRP3 inflammasome protein^54^. These conformers imitate conformational continuum of the NACHT domain rotating around the LRR domain over an angular range of 90° during inflammasome activation^54^ (Fig. 1c). The particles for each conformer are uniformly distributed in each dataset (see Methods). We conducted blind assessments on 3D classification and heterogeneous reconstructions by AlphaCryo4D, without providing any information of particle orientations, translations and conformational identities (Fig. 1d-f, Supplementary Fig. 2, Methods). The 3D classification precision of a retrieved conformer was computed as the ratio of the particle number of correct class assignment (based on the ground truth) versus the total particle number in the class. The results were then compared with several alternative methods, including conventional maximum-likelihood-based 3D (ML3D) classification in RELION^3, 4, 6^, 3D variability analysis (3DVA) in cryoSPARC^2, 11^, and deep generative model-based cryoDRGN^12^. In all blind tests by presetting the class number to 20, AlphaCryo4D retrieved all 20 conformers and markedly outperformed the alternative methods, with an average of 3D classification precision at 0.83, 0.82 and 0.65 for datasets with SNRs of 0.05, 0.01 and 0.005, respectively (Fig. 2a, Supplementary Figs. 3a, 4). By contrast, all alternative methods missed two to seven conformers entirely and exhibited 3D classification precisions in the range of 0.2-0.5 in general (Fig. 2b-d, Supplementary Fig. 3c-k). By increasing the preset class numbers to 25 or 30 that is more than the number of ground-truth conformers, all tested methods appear to be improved in the precision of 3D classification marginally but also reduced in the recall of classification (defined as the percentage of correctly assigned particles versus the number of ground-truth particles for a conformer) (Supplementary Fig. 5). In these cases, AlphaCryo4D still outperformed all alternative methods considerably, with the highest average classification precision reaching 0.91 at the SNR of 0.05.

**Fig. 2.**
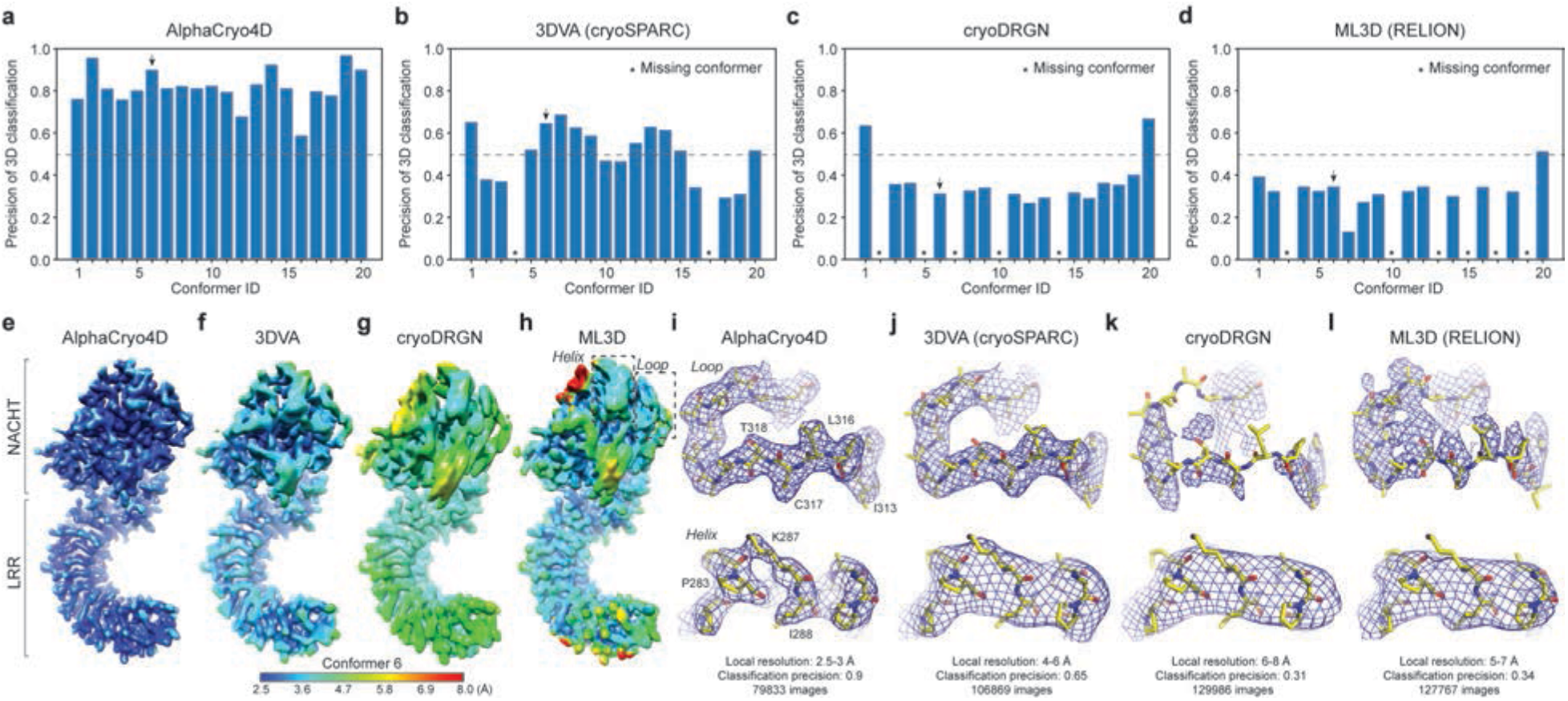
Performance evaluation of AlphaCryo4D for reconstructions of conformational continuum at the atomic level. **a**-**d,** Plots of 3D classification precisions of the 20 NLRP3 conformers from blind assessments on the simulated NLRP3 dataset with SNR of 0.01, using AlphaCyo4D (**a**), 3DVA algorithm in cryoSPARC^11^ (**b**), cryoDRGN^12^ (**c**), and conventional ML3D in RELION^3^ (**d**). All 3D conformers in panel (**a**) were reconstructed to 2.6-2.9 Å resolution (Supplementary Fig. 3l). Asterisks mark the missing conformers that were completely lost due to misclassification by 3DVA, cryoDRGN and ML3D. **e**-**h**, Typical side-by-side comparison of density map quality and local resolution of the same conformer (ID 6) reconstructed by AlphaCryo4D (**e**), 3DVA (**f**), cryoDRGN (**g**) and ML3D (**h**). The maps are colored by their local resolutions calculated by Bsoft blocres program. **i**-**l,** Closeup side-by-side comparison of the same two secondary structures, including a loop (upper row) and a helix (lower row), in the NACHT domain illustrates considerable improvements in local density quality and resolution by AlphaCryo4D (**i**) as opposed to 3DVA/cryoSPARC (**j**), cryoDRGN (**k**) and ML3D/RELION (**l**). The locations of the loop and helix in the NLRP3 structure are marked by dashed boxes in panel (**h**). The same ground-truth atomic model of Conformer 6 shown in stick representations is superimposed with the density maps shown in blue mesh representations, all from the same perspective. The atomic model was not further refined against each map for visual validation of the map accuracy.

The 3D classification precision appears to be strongly correlated with the map quality and resolution homogeneity across the density map (Fig. 2, Supplementary Fig. 2h). All density maps from AlphaCryo4D consistently show homogeneous local resolutions (at 2.6-2.9 Å for SNR of 0.01) between the NACHT and LRR domains (Fig. 2e, i, Supplementary Fig. 2h, 3l, 4a). By contrast, all density maps by the alternative methods show lower average resolutions and notably heterogeneous local resolutions, with the NACHT domain exhibiting substantially lower resolution than that of the LRR domain, causing blurred features, broken loops and invisible sidechains in NACHT (Fig. 2f-h, j-l, Supplementary Figs. 3m-o, 4b-d). Thus, the significantly improved 3D classification accuracy by AlphaCryo4D enables 4D reconstruction of conformational continuum at the atomic level.

Having tested AlphaCryo4D on the simulated datasets with evenly distributed conformers, we further generated a simulated dataset of 0.01 SNR with a non-uniform distribution of the 20 conformers that follows a triangular wave function and presents a more challenging scenario (Supplementary Fig. 6a). Application of AlphaCryo4D on this dataset exhibited only slight reduction in 3D classification accuracy compared to the results on the datasets of evenly distributed conformers, and it still outperformed other methods tested on the uniformly distributed datasets. Moreover, the particle distribution computed by AlphaCryo4D appears to approximately recapitulate the triangular wave function (Supplementary Fig. 6b-f), suggesting that AlphaCryo4D is potentially capable of reconstituting the overall statistical distribution of underlying conformational continuum.

To understand how the 3D classification accuracy is improved, we further studied the algorithmic mechanism by ablating or replacing certain components of AlphaCryo4D in 24 control tests (Supplementary Figs. 2d-g, 7, see Methods for details). By removing the component of deep manifold learning from AlphaCryo4D, the statistical distributions of high-precision 3D classes after particle voting are reduced by ∼60% (Supplementary Fig. 7d-f). Similarly, classifying particles without particle voting or in the absence of deep residual autoencoder reduces the high-precision 3D classes by ∼15-30% at a lower SNR (Supplementary Fig. 7m-r).

Further, replacing t-SNE with other four dimensionality reduction techniques resulted in the loss of at least half of ground-truth conformers in 3D classification, including PCA, multidimensional scaling (MDS)^55^, isometric mapping^56^ and locally linear embedding^57^ (Supplementary Fig. 2d-g). Throughout AlphaCryo4D, the steps of particle shuffling, defining voting boundaries on energy landscapes and energy-based particle voting contribute to ∼16%, ∼20% and ∼40% improvements of high-precision 3D classes, respectively (Supplementary Fig. 7g-l, s-u). Taken together, these results indicate that all components of AlphaCryo4D contribute to the improved accuracy of 3D classification.

### Breaking the resolution limit of the dynamic domains of RyR1

Having conducted the proof-of-principle study of AlphaCryo4D using the simulated datasets, we then turned to five experimental datasets. We first applied AlphaCryo4D to solve the dynamic domains of the 2.2-MDa ryanodine receptor (RyR), a tetrameric calcium-activated channel protein complex that mediates the release of calcium ions from intracellular stores. The mammalian RyR1 has been determined at near-atomic resolution in its central core in multiple conformations^58, 59^, which was recently further improved to sub-3 Å resolution^45^. However, approximately half of the entire RyR1 structure located at peripheral domains remain at moderate to low resolution due to their complex dynamics that precludes atomic modeling with reliable amino acids assignment. Extensive previous efforts have failed in further improving the resolution of the peripheral domains by the conventional methods^45, 58, 59^. Thus, we applied AlphaCryo4D to analyze the RyR1 dataset^45^ to test if our approach can overcome the conformational heterogeneity and break the resolution barrier. We computed a focused energy landscape using 310 bootstrapped density maps to characterize the conformational distribution of the peripheral domains (see Methods). The energy landscape reveals three new sub-states from the original dataset corresponding to state 1 conformation^45^ (Fig. 3a, Supplementary Fig. 8a). The peripheral domains of the three states vibrate vertically against the central domains with a very small amplitude (Fig. 3b, Supplementary Fig. 8b), which is sufficient to blur the local density when the entire particles are used for image alignment.

**Fig. 3.**
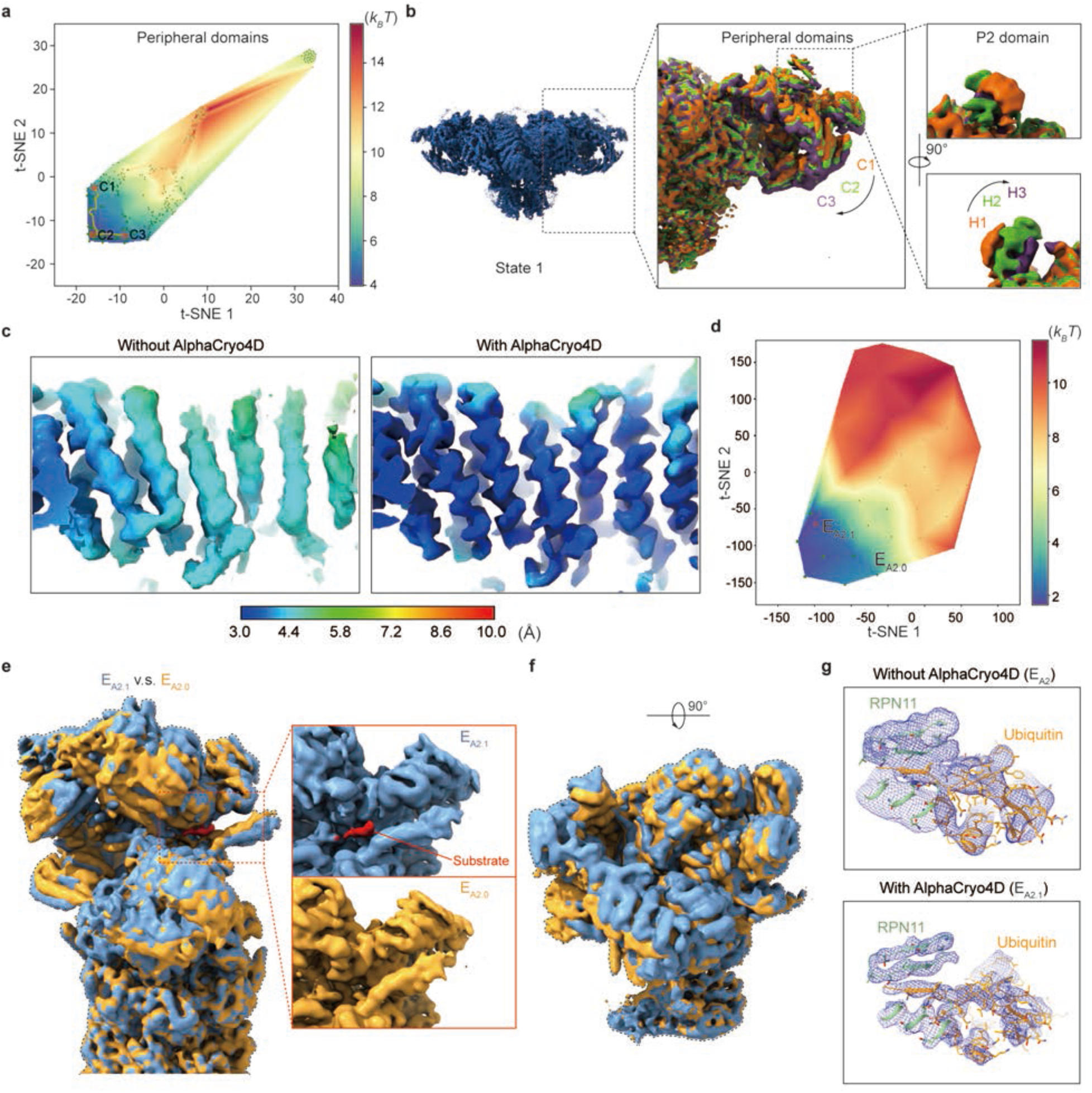
Resolution improvement of the dynamic components of RyR1 and the 26S proteasome by AlphaCryo4D. **a,** Energy landscape of the peripheral domains of the RyR1. Clusters C1, C2 and C3 were selected for particle voting by this energy landscape. **b,** Cryo-EM density maps of clusters C1-C3 and H1-H3 classified by AlphaCryo4D. The peripheral domains exhibited hierarchical dynamics in its P2 domain, which was not well observed in the original density map of state 1. **c,** Comparison of the helical domain density map of the original state 1without and with AlphaCryo4D. The process described here improved the resolution of the density. **d,** Energy landscape calculated on the regulatory particle of the proteasome state E_A2_. Clusters E_A2.0_ and E_A2.1_ were determined on this energy landscape, revealing the states of substrate-unbound and substrate-bound RPN11, respectively. **e** and **f,** Cryo-EM density map of cluster E_A2.0_ superimposed with that of cluster E_A2.1_ from two orthogonal viewing angles. Side view, panel (**e**). Top view, panel (**f**). The lid subcomplex undergoes conformational changes during substrate binding. Insert of panel (**e**) shows closeup side-by-side comparison of the two states. The red region denoted the substrate in the density map of 26S proteasome. **g,** Quality and resolution improvement of local cryo-EM density map of the RPN11-bound 8.5-kDa ubiquitin in state E_A2.1_ (lower panel) relative to the previously published structure^23^ (upper panel).

Using the AlphaCryo4D-classified particles for refinement, the local resolution of the peripheral domains was refined to 3.3 Å, representing an unprecedented improvement over the previous resolution of 5-8 Å for nearly half of RyR1. The resulting density map showed high-resolution features including helical pitches and side-chain densities (Fig. 3c, Supplementary Fig. 8f). However, the small P2/RY3&4 domain abutting the helical C-Sol domain via flexible links still has low resolution in the refined map. To improve the visualization of P2, we computed the zoomed-in energy landscape by focusing on the P2 domain. This energy landscape also exhibited three conformational clusters (Supplementary Fig. 8e), from which we found the P2 domain oscillating relative to the helical C-Sol domain. Further refinement of the AlphaCryo4D-classified particles improved the clarity of the secondary structure in the density map, but it did not reach the atomic level likely due to the very small size of this domain limiting the alignment accuracy (Supplementary Fig. 8c). In summary, the application to RyR1 demonstrates that AlphaCryo4D is capable of breaking the resolution barrier of the dynamic domain by resolving hierarchically entangled conformational dynamics of the RyR1 peripheral domains.

### Visualizing hidden dynamics of the 26S proteasome

The 26S proteasome is one of the most complex, dynamic, heterogeneous holoenzyme machineries that regulates nearly all aspects of cell physiology in eukaryotes^23, 60–63^. Visualizing atomic structures of nonequilibrium intermediates connecting the major states of the functional proteasome in the act of substrate degradation has been unsuccessful^4, 6, 7, 23, 60^. To test this possibility, we applied AlphaCryo4D to analyze a large experimental dataset of substrate-engaged human 26S proteasome (EMPIAR-10669)^24^, which comprises more than 3 million particles. AlphaCryo4D was able to significantly enrich the number of visualized proteasome conformers to 64 using this dataset, which was previously used to determine atomic structures of seven proteasome states^23^. As the majority of the states were refined to near-atomic resolution, they allowed us to obtain a more complete picture of the functional dynamics of the proteasome at the atomic level.

To demonstrate that AlpahCryo4D can simultaneously improve 3D classification and resolution of key dynamic features, we examine a representative case among numerous improvements of solving the proteasome dynamics by AlphaCryo4D^24^. Here we focus on state E_A2_ of the substrate-engaged 26S proteasome, which is the initial state that a substrate-conjugated ubiquitin moiety binds the deubiquitylating enzyme RPN11 to prepare for substrate engagement and deubiquitylation. In this case, we expected to use AlphaCryo4D to detect if an intermediate substrate binding to RPN11 proceeds the deubiquitylation step of proteasome^23^. Previous reconstruction on state E_A2_ has showed that no substrate but ubiquitin at a moderate resolution was bound to the RPN11 surface^23^. On the energy landscape of the sub-dataset corresponding to state E_A2_, we examined two conformational clusters, designated E_A2.0_ and E_A2.1_, which were refined to 3.5 and 3.2 Å, respectively (Fig. 3d). Comparing these two density maps reveals subtle conformation changes of the lid subcomplex against the base, although both show no substrate binding in the central channel of hexameric AAA-ATPase ring (Fig. 3e). Markedly, state E_A2.1_ exhibits a segment of substrate bound to the cryptic hydrophobic groove at the interface between RPN11 and RPT4 near the active site of RPN11, whereas no substrate is observed at the same site in state E_A2.0_ (Fig. 3f). This polypeptide substrate segment has a length of about 4 amino acids, which is comparable in size to many small-molecule compounds, suggesting that AlphaCryo4D may be capable of classifying cryo-EM data towards very small features relevant to drug discovery.

The conformation of state E_A2.0_ appears identical to that of the previously published state E_A2_. The resolution of the RPN11-bound, 8.5-kDa ubiquitin density in state E_A2_ is around 6-9 Å, although the overall resolution of state E_A2_ was measured to 3.3 Å. Approximately half particles previously classified to E_A2_ were also classified to E_A2.1_, indicating the mixture of different conformers that caused the low-resolution feature of ubiquitin in previous studies^23^. Notably, this RPN11-bound ubiquitin in E_A2.1_ clearly shows higher resolution features consistent with 3.8 Å, with sufficiently separated β-strands (Fig. 3g). It indicates that the RPT5 N-loop, C-terminal strand of ubiquitin and insert-1 β-hairpin of RPN11 already form a four-stranded β-sheet prior to substrate insertion into the proteasomal AAA-ATPase channel. These features were only sufficiently observed in state E_B_ previously, where the substrate has completed its insertion into the AAA-ATPase ring^23^. These high-resolution features allow us to confidently interpret state E_A2.1_ as a transient intermediate between the previously published states E_A2_ and E_B_. Altogether, our extensive application of AlphaCryo4D to the experimental dataset demonstrates its capability in exploring the ‘hidden’ conformational space of substrate-engaged proteasome at the atomic level, thus providing unprecedented insights into the molecular mechanism of proteasome autoregulation as described in detail elsewhere^24^.

### Revealing residual conformational heterogeneity in the *Pf*80S ribosome

We further evaluated the capability of AlphaCryo4D in detecting residual heterogeneity in a mostly homogeneous dataset of the Plasmodium falciparum 80S (*Pf*80S) ribosome bound to anti-protozoan drug emetine (EMPIAR-10028)^46^. The original cryo-EM analysis of the *Pf*80S ribosome dataset (105,247 particles) determined a major conformational state and detected flexibility in the head region of the 40S small subunit, which was originally unsolved^46^. Application of AlphaCryo4D to analyze this dataset reveals notable rotation of 40S in the *Pf*80S ribosome. By reconstitution of the approximately continuous energy landscape using 79 density maps, we simultaneously identified two 40S-rotated states (R3 and R4) and another state (R2) with missing 40S head in addition to the original conformational state (R1) located in the deepest energy well (Fig. 4a, b). Notably, AlphaCryo4D was able to maintain the original resolution of the major state at 3.2 Å, while allowing the three new states R2, R3 and R4 to be refined to 3.6, 4.1 and 4.6 Å, respectively, without using more data (Supplementary Fig. 9).

**Fig. 4.**
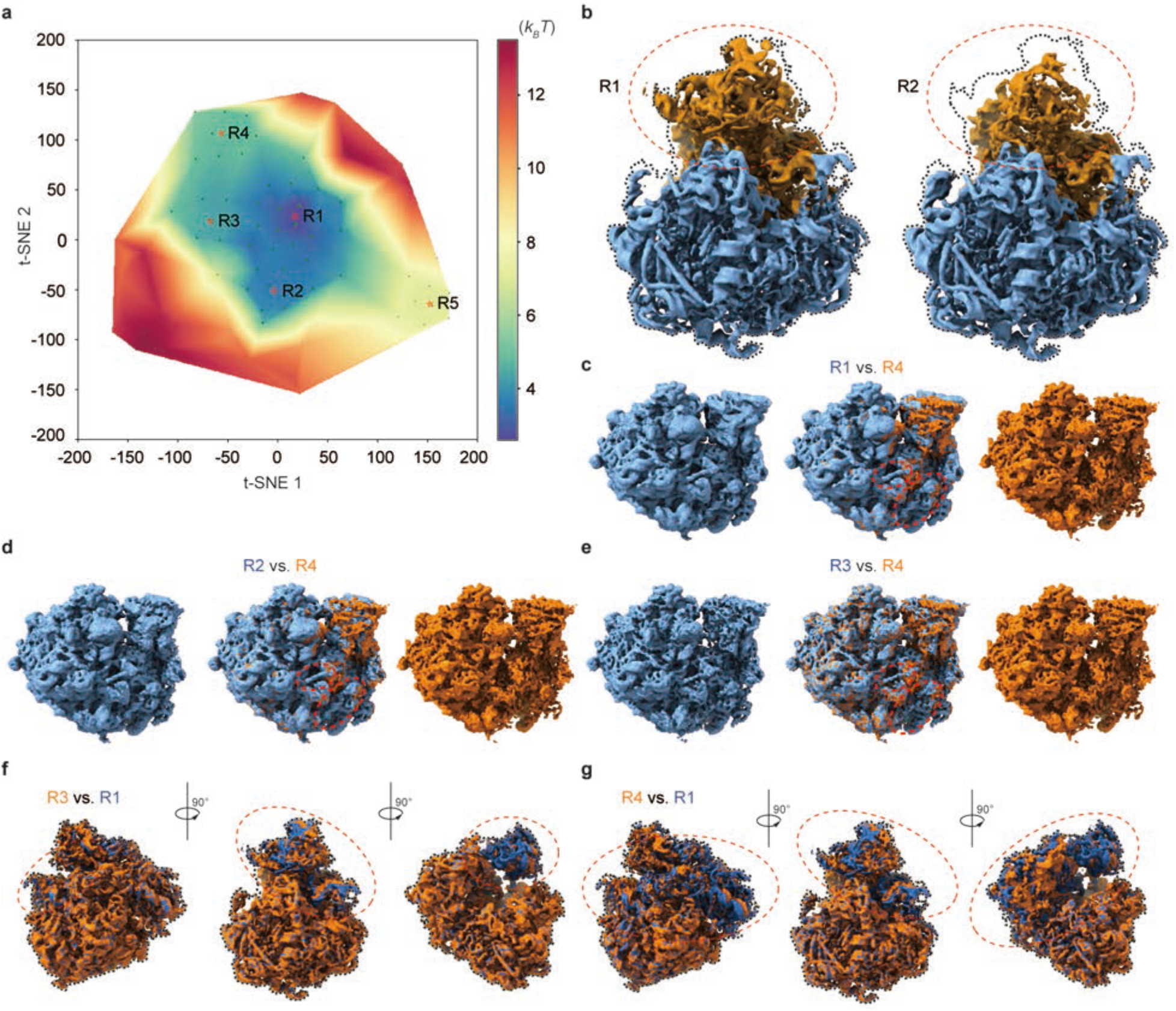
Visualizing conformational dynamics of the *Pf*80S ribosome bound to anti-protozoan drug at high resolution by AlphaCryo4D. **a,** Clusters R1-R5 on the energy landscape by AlphaCryo4D. Energy-based particle voting were conducted within each cluster to keep particles of high reproducibility. **b,** Comparison of the density maps of clusters R1 and R2. Cluster R1 is the major state in the *Pf*80S ribosome dataset, while cluster R2 accounts for the state missing the head of 40S (red circles). The dynamic region of the density map is colored orange and the stable region is colored blue. **c**-**e,** Comparisons of states R1 and R4 (**c**), states R2 and R4 (**d**), and states R3 and R4 (**e**) reveal the differential appearance of the C-terminal helix of eL8 and an rRNA helix. Note that the rRNA helix is only partially presented in state R3. **f,** State R3 superimposed with state R1 from three different viewing angles. The density of the 40S head region (red circles) rotated relative to the 60S. **g,** State R4 superimposed with state R1 from three different viewing angles. The density of the whole 40S (red circles) is rotated relative to the 60S.

On the energy landscape, the distance between states R4 and R1 is longer than that between states R3 and R1, indicating that a larger conformational change occurs in state R4. Indeed, superimposing the density maps of states R3 and R1 together reveals that the rotation of 40S is localized within its head region with concerted movement in the central protuberance of 60S, whereas the rest of the 40S exhibits no obvious movement (Fig. 4f). By contrast, the entire 40S rotates above the 60S in state R4 relative to R1, and this coincides with the intrinsic structural rearrangement within 60S near the central protuberance and L1 stalk, which appears to also drive the 40S translation relative to the 60S (Fig. 4g), indicating a different mode of intersubunit rotation. Moreover, the reconstructions of R3 and R4 also identified differential rearrangements of small structural elements, including the stepwise disappearance of an rRNA helix and of the intersubunit bridge formed by eL8 C-terminal helix, in line with previous studies on the *Pf*80S dynamics during translation^64^ (Fig. 4c-e). Our analysis suggests that the superficially simple rotation of the 40S subunit is a manifestation of complicated conformational dynamics involving the entire *Pf*80S ribosome. Given that the *Pf*80S ribosome was bound to the anti-protozoan drug visible in all new states (Supplementary Fig. 9e), the AlphaCryo4D-enabled analysis of structural dynamics at high resolution opens the door to investigate the drug-modulated allosteric effect relevant to therapeutic improvement^46^.

### Visualizing conformational continuum of the pre-catalytic spliceosome

We next evaluated the applicability of AlphaCryo4D in simultaneously visualizing continuous motion and discrete compositional heterogeneity in the pre-catalytic spliceosome (EMPIAR-10180)^47^. Previous study using focused classification has provided a multistate-averaged map of the pre-catalytic spliceosome and suggested that the SF3b subcomplex may sample a continuum of conformations. To characterize the pre-catalytic spliceosome dynamics, we bootstrapped 160 density maps using 327,490 particles to reconstitute its energy landscape (Fig. 5a), which allowed us to determine many intermediate states (S1-S9) representative of a continuous motion of the SF3b and helicase subcomplexes as well as several compositionally distinct states (S10-S14). On the energy landscape, the nine states S1-S9 representing the continuous motions are intuitively located in adjacent clusters, forming a major MEP traversing the long axis of the energy landscape (Fig. 5c). These states show both independent movements of the SF3b and helicase, as well as concerted motion of the two subcomplexes (Supplementary Fig. 10a). For instance, comparison between states S3 and S4 exhibits that the SF3b and helicase move away from each other, whereas states S5 and S6 show co-directional motion of both the SF3b and helicase. Interestingly, the Cus1 subunit in the SF3b subcomplex appears to switch its contact between the N-terminal helicase cassettes of RecA-1 and RecA-2, implicating that part of conformational rearrangement is mediated by the transient interface B between the SF3b and helicase^47^.

**Fig. 5.**
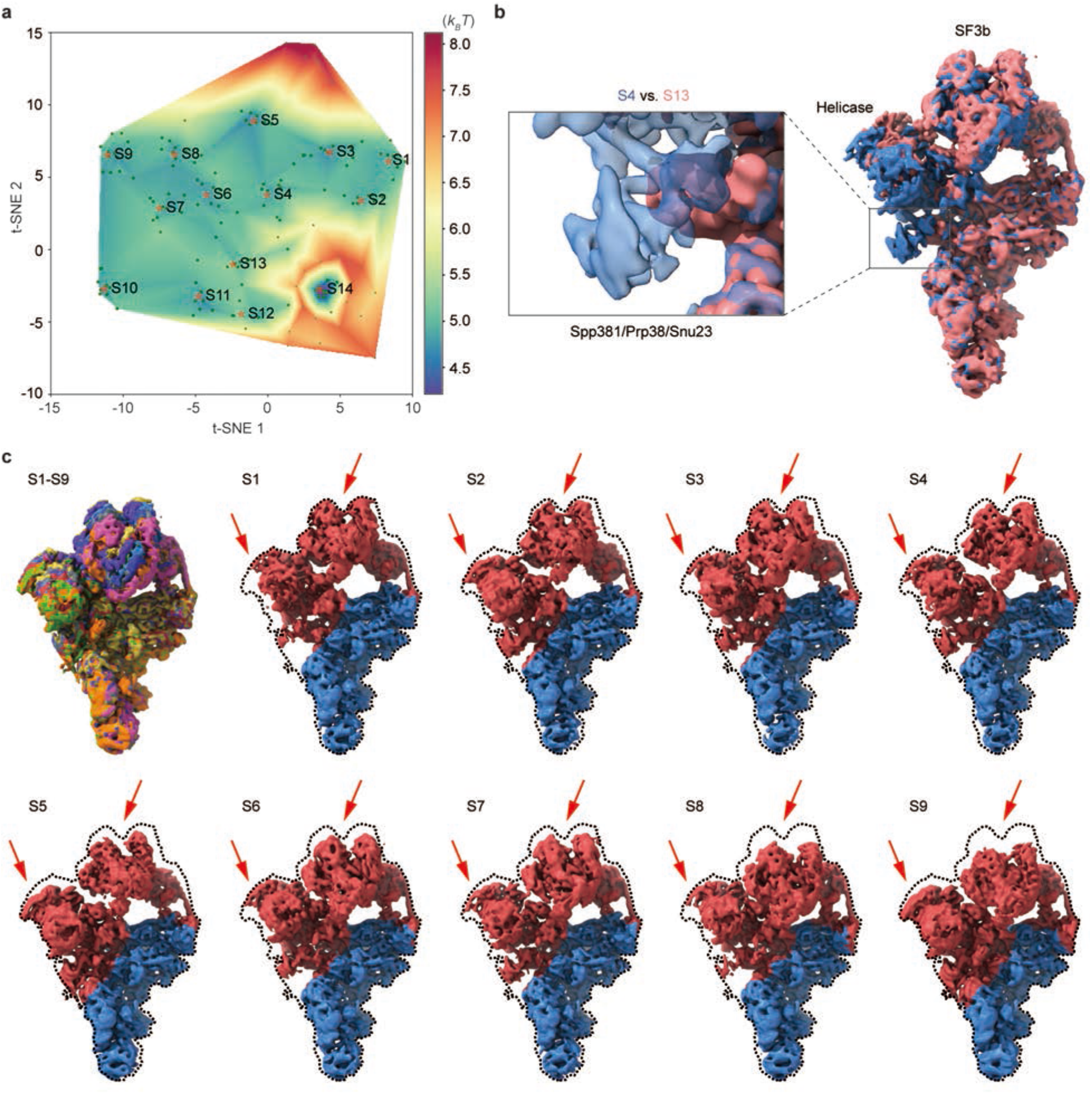
Choreographing continuous inter-subunit motions of the pre-catalytic spliceosome by AlphaCryo4D. **a,** Energy landscape by AlphaCryo4D uncovering 14 clusters designated as S1-S14. Clusters S1-S9 are related to the continuous rotational motion of the SF3b and helicase subcomplexes of pre-catalytic spliceosome, and the density maps of clusters S10-S14 are shown in Supplementary Fig. 10b. **b,** Density map of the Spp381/Prp38/Snu23 missing state S13 superimposed with the state S4. Closeup comparison showed the density difference between states S13 and S4. **c,** Cryo-EM density maps of clusters S1-S9. The dynamic component of the density map was colored red and the static component was colored blue. The red arrows suggested the rotational motion of the SF3b and helicase subcomplexes within the dashed outline.

In addition to mapping the continuous inter-subunit motion, AlphaCryo4D also reveals potential allosteric effect of local structural variation relevant to spliceosome activation. Notably, comparison between states S4 and S13 exhibits that the U2 snRNP and helicase motions among these states appear to be allosterically coupled to the disappearance of the Spp381, Prp38 and Snu23 densities in state S13 (Fig. 5b). As the proteins Spp381, Prp38 and Snu23 are required for spliceosome activation^47^, these coexisting conformers may be important to understand the intermediate steps of spliceosome activation. These biologically important dynamic features have not been previously reported from the same dataset. Taken together, our analysis demonstrates that AlpahCryo4D is capable of characterizing continuous conformational changes on a dataset with a moderate size.

### Discovery of hidden assembly intermediates of bacterial ribosome

In the last application, we used AlphaCryo4D to analyze a highly heterogeneous dataset of the *Escherichia coli* 50S large ribosomal subunit (LSU) (EMPIAR-10076)^48^. This dataset (131,899 particles) is known to be both compositionally heterogeneous and conformationally dynamic in that the 50S LSU undergoes bL17-depleted intermediate assembly. We first computed the overall energy landscape of the ribosomal assembly using 119 density maps (Fig. 6b). On this energy landscape, the originally reported five major states A-E were straightforwardly reproduced by energy-based particle voting. Obviously, state A containing both 50S and 30S subcomplexes is a rare state distinct from other conformers (Fig. 6a). The density map of state A reconstructed by only 766 particles still retains enough details, benefiting from the strategy of energy-based particle voting.

**Fig. 6.**
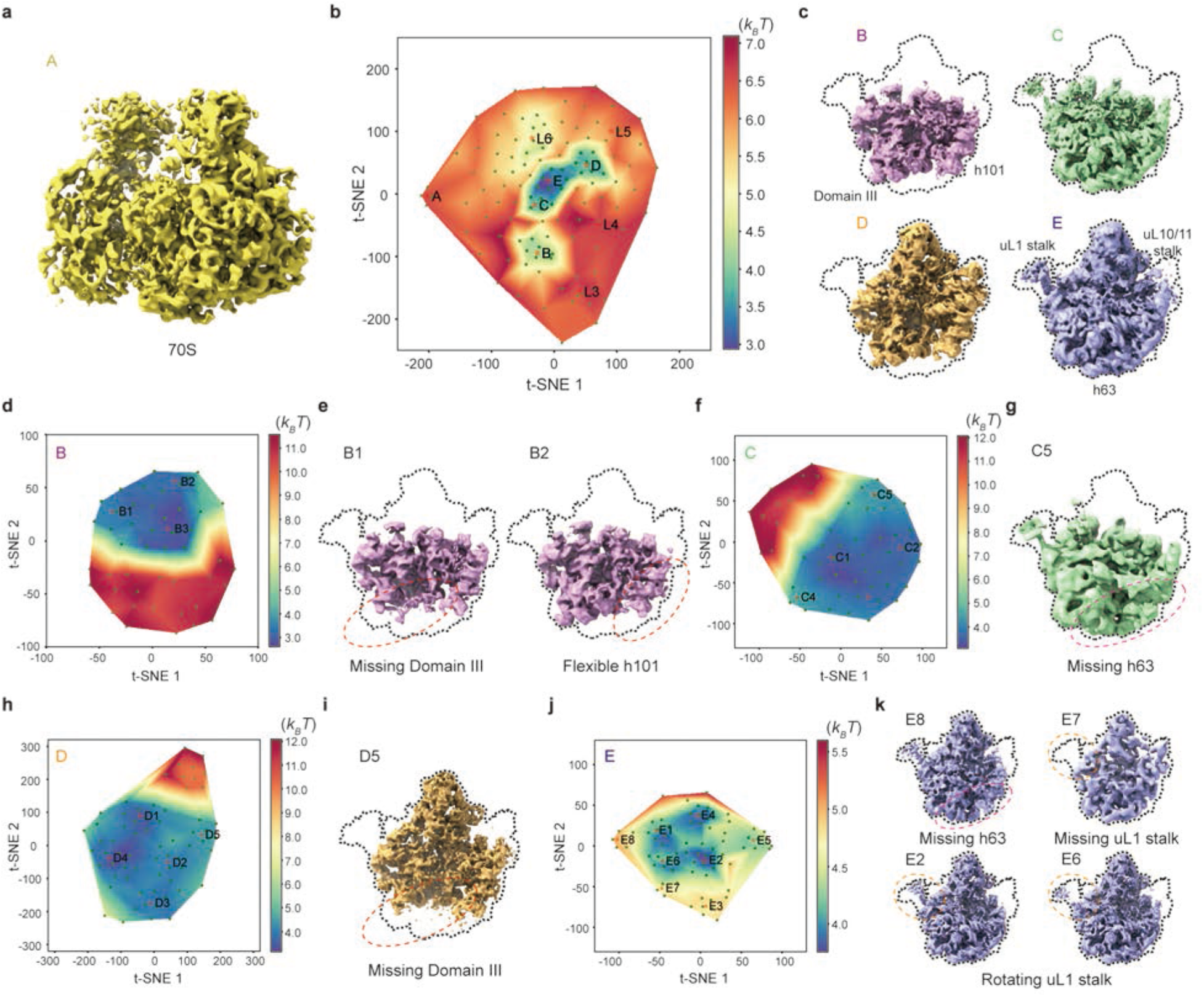
Exploring hidden conformational space of the bacterial ribosomal assembly intermediates by AlphaCryo4D. **a,** Cryo-EM density map of state A determined by AlphaCryo4D. This state was located far from other states on the energy landscape shown in (**b**). **b,** Global energy landscape of bacterial ribosomal assembly intermediates. Clusters A-E and L3-L6 were reconstructed for further analysis. **c,** Density maps of the major states B, C, D and E reconstructed by AlphaCryo4D. The density maps of these four states were refined using the particles of clusters B, C, D and E respectively from the energy landscape. **d,** Zoomed-in energy landscape of the major state B. **e,** New subclass conformers B1 and B2 of state B. Comparing to the original state B (Supplementary Fig. 11b), the new states B1 and B2 appear to miss Domain III and show flexible rRNA helix 101, respectively, as discovered by AlphaCryo4D. **f,** Zoomed-in energy landscape of the major state C generating clusters C1-C5. **g,** A new state (C5) of C determined by AlphaCryo4D. The rRNA helix 63 was not observed in the density map of C5. **h,** Clusters D1-D5 from the zoom-in energy landscape of the major state D. **i,** A newly discovered minor state D5. Like the minor state B1 of the major state B, Domain III was not seen in the density map of D5. **j,** Clusters E1-E8 on the zoom-in energy landscape of the major state E by AlphaCryo4D. **k,** Discovery of new subclass conformers in the major state E. Like C5, the new state E8 lacks the rRNA helix 63. Comparing to the previously reported state E3 (Supplementary Fig. 11e), the uL1 stalk is not found in the subclass E7. Two new states, E2 and E6, with different conformational changes of the uL1 stalk were discovered by AlphaCryo4D.

As the remaining particle numbers of states B, C, D and E are all adequate for deeper classification, we then computed their zoomed-in energy landscapes separately. This allowed us to discover seven new conformational states (designated B1, B2, C5, D5, E6, E7, E8) that have evaded all other methods^11, 12, 19, 48^ (Fig. 6d-k), as well as all 13 previously reported sub-class conformers classified from this dataset^12, 48^ (Supplementary Fig. 11). On the zoomed-in energy landscape of state B, state B1 exhibits a low occupancy of domain III as well as another new state D5 that is perhaps a least mature LSU intermediate on the assembly pathway (Fig. 6e, i). The rRNA helix 101 exhibits a small rotation between states B2 and B3 (Fig. 6e and Supplementary Fig. 11b). The differential occupancy of rRNA helix 63 and the movement of uL1 stalk are observed in states C5 and E6-E8 (Fig. 6g, k). Among the sub-states derived from state E, the new state (E7) with uL1 missing was reconstructed by only 357 particles (Fig. 6k), which was another rare state discovered via AlphaCryo4D that is only 0.3% of the entire dataset. These results suggest that AlphaCryo4D is highly efficient in finding transient states of extremely low populations and in exploring the ‘dark’ conformers in the conformational space of highly heterogeneous systems.

## Discussion

One primary objective of the AlphaCryo4D development is to simultaneously improve reconstruction resolution and 3D classification without assumption on the behavior of sample dynamics and heterogeneity. A key lesson from this study is that the improved accuracy of 3D classification is often positively correlated with the resolution gain in general, as misclassification is presumably a key factor limiting achievable resolution. Because of extremely low SNR, most objective functions in image similarity measurement and machine learning algorithms tend to poorly perform or entirely fail on cryo-EM data. To date, no methods have included any implicit metrics or strategies for validating 3D classification accuracy in cryo-EM. We address this issue by introducing *M*-fold particle shuffling method and the energy-based particle voting algorithm in AlphaCryo4D that automatically checks the reproducibility and robustness of the 3D classification. This allows users to objectively reject non-reproducible particles and permits a maximal number of particles to be assessed and classified in an integrative procedure based on uniform, objective criteria. Our extensive tests suggest that the particle-voting algorithm synergistically enhances the performance of deep manifold learning. Moreover, it can potentially rescue certain particles that are prone to be misclassified if processed only once by deep manifold learning, thus boosting the efficiency of particle usage without necessarily sacrificing quality and homogeneity of selected particles. This mechanism simultaneously optimizes the usage of all available particles and their 3D classification accuracy for achieving higher resolution.

Our particle-shuffling and Bayesian bootstrap strategy is conceptually related to the stochastic bootstrap method previously proposed for estimating 3D variance map^65^ or performing 3D PCA ^13^ to detect conformational variability. One essential difference lies in that our bootstrap requires that only particles of similar or identical conformations via likelihood-based similarity estimation are grouped together instead of being combined or resampled in a stochastic manner^1, 18^. Another difference is that the goal of *M*-fold particle shuffling is to improve the quality of reconstituted energy landscape and to enable energy-based particle voting for intrinsic cross-validation of 3D classification. Furthermore, AlphaCryo4D does not assume any prior knowledge regarding the conformational landscape, its continuity and topology, as well as chemical composition of the macromolecular system. Our approach allows multiple consensus models being used for optimizing the particle alignment during Bayesian bootstrap. This can theoretically deal with more complicated conformational dynamics when no stable core structure is available to guide the consensus alignment over a single model.

Using energy landscape to represent the conformational variation provides several advantages. First, it has theoretical roots in physical chemistry and protein dynamics, in which extensive research tools being built around the energy landscape and transition-state theory allows correlation with other complementary approaches, such as single-molecule florescence microscopy, NMR and molecular dynamics simulation. Second, the energy landscape allows users to intuitively examine kinetic relationships between the adjacent conformers and to potentially discover new conformations. Our approach to use many bootstrapped density maps to estimate the ideologically continuous energy landscape is statistically analogous to other methods proposed to map single particles to continuous distributions in the latent space^11, 12, 19^. The latter may suffer from the lack of energetic information that can restrict the interpretation of the resulting reconstruction. Third, while our method of mapping 3D volumes to manifolds increases the sampling grain in approximating the continuity of conformational distribution as compared to other methods directly mapping single particles to the latent space^11, 12, 19^, AlphaCryo4D trades off the fineness of sampling grain in the continuity approximation for resolution gain. In nearly all cases we examined, AlphaCryo4D apparently still oversampled the conformational space at large in terms of finding those conformational states that can potentially achieve high resolution.

The limitation of AlphaCryo4D lies in that it requires generally larger datasets and more computational costs to fully exploit its advantages and potentials. The data size must proportionally scale with and match the degree of conformational heterogeneity in order to visualize a full spectrum of conformational states at the atomic level. Moreover, its outcomes depend on the success of initial consensus alignment of all particles during particle shuffling and volume bootstrapping. Certain conformational dynamics or heterogeneity can interfere with image alignment, in which case the performance of AlpahCryo4D will be restricted by the alignment errors in during consensus refinement. This is a common problem for all existing methods developed so far. Our tentative solution is to use multiple consensus models for improving image alignment. Failure in obtaining accurate alignment parameters can lead to futile, erroneous estimation of conformational heterogeneity in the subsequent steps no matter which methods are used.

The use of t-SNE to reconstitute the energy landscape poses both potential advantages and caveats. The t-SNE algorithm can well preserve the local topology but is not warranted to preserve global topology, which is nonetheless a common limitation for existing manifold learning techniques. This issue is partly alleviated by the particle-voting algorithm that is performed at the local areas of energy minima on the energy landscape. Dimensionality reduction to 2D by t-SNE, as well as MEP searching on a 2D energy landscape might also limit its capability in disentangling complex dynamics. Further investigations are required to understand how to map individual particles to a continuous energy landscape without trading off reconstruction resolution, to estimate the energy landscape at higher dimensions, to preserve its global topology with high fidelity and to fully automatize MEP solution.

## Methods

### *M*-fold particle shuffling and Bayesian bootstrapping

A key philosophy of the AlphaCryo4D design is to avoid subjective judgement on the particle quality and usability if it is not apparent false particles like ice contaminants, carbon edges and other obvious impurities. Deep-learning-based particle picking in DeepEM^66^, Torpaz^67^ or other similarly performed software is favored for data preprocessing prior to AlphaCryo4D. To prepare particle datasets for AlphaCryo4D, an initial unsupervised 2D image classification and particle selection, preferentially conducted by the statistical manifold-learning-based algorithm in ROME^5^, is necessary to ensure that no apparent false particles are selected for further analysis and that the data have been collected under an optimal microscope alignment condition, such as optimized coma-free alignment. No particles should be discarded based on their structural appearance during this step if they are not apparent false particles. Any additional 3D classification should be avoided to pre-maturely reject particles prior to particle shuffling and volume bootstrapping in the first step of AlphaCryo4D processing. Pre-maturely rejecting true particles via any 2D and 3D classification is expected to introduce subjective bias and to impair the native conformational continuity and statistical integrity intrinsically existing in the dataset.

In raw cryo-EM data, 2D transmission images of biological macromolecules suffer from extremely heavy background noise, due to the use of low electron dose to avoid radiation damage. To tackle the conformational heterogeneity of the macromolecule sample of interest in the presence of heavy image noise, the particle shuffling and volume bootstrapping procedure was devised to incorporate the Bayesian or maximum-likelihood-based 3D clustering in RELION^3^. To enable the particle-voting algorithm in the late stage of AlphaCryo4D, each particle is reused *M* times during particle shuffling to bootstrap a large number of 3D volumes (Supplementary Fig. 1a). First, all particle images are aligned to the same frame of reference in a consensus 3D reconstruction and refinement in RELION^3^ or ROME^5^ to obtain the initial alignment parameters of three Euler angles and two translational shifts. Optimization for alignment accuracy should be pursued to avoid the error propagation to the subsequent steps in AlphaCryo4D. For a dataset with both compositional and conformational heterogeneity, coarsely classifying the dataset to a few 3D classes during initial image alignment, limiting the 3D alignment to a moderate resolution like 10 Å or 15 Å during global orientational search, progressing to small enough angular steps in the final stage of consensus refinement, may be optionally practiced to optimize the initial 3D alignment. Failure of initial alignment of particles would lead to failure of all subsequent AlphaCryo4D analysis.

Next, based on the results of consensus alignment, in the particle-shuffling step, all particles were divided into *M* + 1 groups, and *M* was set to an odd number of at least 3. Then the whole dataset was shuffled *M* + 1 times by removing a different group out of the dataset each time. Each shuffled dataset is classified into a large number (*B*) of 3D volumes, often tens to hundreds, by the maximum-likelihood 3D classification algorithm without further image alignment in RELION (with ‘skip-align’ option turned on). This is necessary because the alignment accuracy often degrades when the sizes of 3D classes decrease considerably. This step is repeated *M* + 1 times, each time on a shuffled dataset missing a different group among the *M* + 1 groups. Due to the effect of *M*-fold particle shuffling, the outcome of this entire process is expected to bootstrap up to thousands of 3D volumes in total (i.e., *B*(*M* +1) > 1000). Each particle is used and contributed to the 3D reconstructions of *M* volumes, which prepare it for the particle-voting algorithm to evaluate the robustness and reproducibility of each particle with respect to the eventual 3D classification.

For processing a large dataset including millions of single-particle images, it becomes infeasible even for a modern high-performance computing system to do the consensus alignment by including all particles once in a single run due to limitation of supercomputer memory and the scalability of the alignment software. To tackle this issue, the whole dataset is randomly split into a number (*D*) of sub-datasets for batch processing, with each sub-dataset including about one to two hundred thousand particles, depending on the scale of available supercomputing system. In this case, the initial reference should be used for the consensus alignment of different sub-datasets to minimize volume alignment errors in the later step. The total number of resulting bootstrapped volumes becomes *BD*(*M* +1) > 1000. This strategy can substantially reduce the supercomputer memory pressure and requirement. In each sub-dataset, all particles were divided into *M* + 1 groups and subject to the particle shuffling and volume bootstrapping procedure as described above.

### Deep residual autoencoder for 3D feature extraction

The bootstrapped volumes may still suffer from reconstruction noises and errors due to variation of particle number, misclassification of conformers and limited alignment accuracy. Thus, we hypothesize that unsupervised deep learning could help capture the key features of structural variations in the bootstrapped volumes and potentially enhance the quality of subsequently reconstituted energy landscapes for improved 3D classification. To this end, a 3D autoencoder was constructed using a deep Fully Convolutional Network (FCN) composed of residual learning blocks^49, 50^. The architecture of the 3D autoencoder consists of the encoder and the decoder, which are denoted as *ε* and 𝒟, respectively (Fig. 1b). The relation between the output ***y*** and the input ***x*** of the network can be expressed as ***y*** = 𝒟(*ε*(***x***)), in which ***x*** is the input 3D density volume with the size of *N*^3^, where *N* is the box size of the density map in pixel units. For reconstruction of the 3D volumes and further optimization, the decoding output ***y*** should be in the same size and range with the input data ***x***. In this way, the framework of FCN is established to restore the input volume, using the sigmoid function 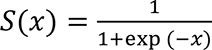 as the activation function of the decoding layer to normalize the value of ***y*** into the range (0, 1). Meanwhile, all 3D density maps ***x*** are normalized with the following equation before input to the deep neural network:

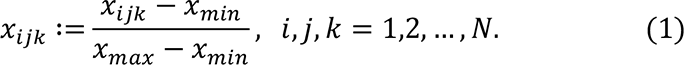

where *x_min_* and *x_max_* are, respectively, the minimum and minimax value in all *x_ijk_*.

The distance between the decoded maps and the input volumes can be used for constructing the loss function to train the 3D kernels and bias of the networks. The value distribution of the encoded 3D feature maps ***z*** = *ε*(***x***) is expected to be an abstract, numerical representation of the underlying structures in the volume data, which may not necessarily have any intuitive real-space physical meanings. The neural network can extract such abstract information in the prediction step, with no restriction on the feature maps ***z*** in the expression of training loss. The loss function is then formulated as:

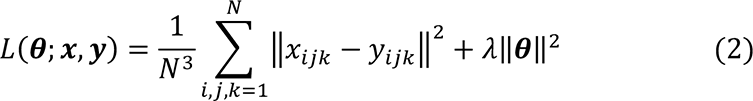

where ***θ*** denotes the weights and bias of the network, and *λ* is L2 norm regularization coefficient. As the feature of the complex structure is difficult to be learned from the 3D volume data, the value of *λ* is set to 0.001 in default to prevent overfitting in the applications to experimental datasets. But it is set to 0 in the tests on the simulated datasets (see below).

To improve the learning capacity of the 3D autoencoder, residual learning blocks containing 3D convolutional and transposed convolutional layers are employed in the encoder and decoder, respectively. In each residual block, a convolutional layer followed by a Batch Normalization^68^ (BN) layer and activation layer appears twice as a basic mapping, which is added with the input to generate the output. The mathematical expression of the *l*th block can be shown as:

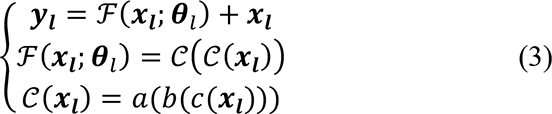

where ***x***_***l***_ and ***y***_***l***_ represent the input and the output of the block, respectively. ℱ(*x*_***l***_; ***θ***_***l***_) denotes the basic mapping of the *l*th block parameterized by ***θ***_***l***_, and 𝒞(***x***_***l***_) is the sequential operation of convolution or transposed convolution *c*, BN *b* and activation *a*. The rectified linear unit (ReLU) function is used in all the activation layers but the last one. In addition, the output of mapping function ℱ(***x***_***l***_; ***θ***_***l***_) must have the same dimension as the input ***x***_***l***_. If this is not the case, the input ***x***_***l***_ must be rescaled along with ℱ(***x***_***l***_; ***θ***_***l***_) using a convolutional or transposed convolutional transformation *c*’(***x***_***l***_) with an appropriate stride value, the parameters of which can be updated in the training step.

### Autoencoder training

To analyze a large number of volumes, we carefully tuned the training of the 3D deep residual autoencoder to obtain suitable kernels and bias. First, parallel computation with multiple GPUs has been implemented to reduce the training time. Then the parameters of the network are optimized by the stochastic gradient descent Adam (Adaptive moment estimation) algorithm, in which the gradients of the objective function *L*(***θ***; ***x***, ***y***) with respect to the parameters ***θ*** can be calculated by the chain rule. Moreover, the learning rate is reduced to one tenth when the loss function does not decrease in three epochs based on the initial value of 0.01. After trained about 50 epochs, the best model is picked to execute the task of structural feature extraction. Using the unsupervised 3D autoencoder, the feature maps ***z*** = *ε*(***x***) encoding the structural discrepancy among the 3D volume data can be extracted automatically without any human intervention.

### Autoencoder hyperparameters

The recommended hyperparameters for the deep residual autoencoder architecture are provided in Supplementary Table 1. Although we expand the residual network (ResNet) into 3D, we keep the original design rules of ResNet for constructing the 3D autoencoder^49, 50^. The first and last convolutional layers have 5 × 5 × 5 filters (kernels). The remaining convolutional layers have 3 × 3 × 3 filters. Because the cubic filter in convolutional layer is computationally expensive, only one 3D filter is used in each of the last three convolutional layers in the encoder and of the first two and last transposed convolutional layers in the decoder. To accommodate a large volume input, two filters are used in the first three convolutional layers and in the third and fourth transposed convolutional layers. Downsampling is directly performed by convolutional layers with a stride of 2. The output dimension of encoding layers is set to be a quarter of the input dimension to avoid over-compressing the feature maps. We employed six 3D convolutional layers in the encoder and five transposed convolutional layers in the decoder based on the expected tradeoff between the learning accuracy and training cost, which is roughly comparable to a 2D ResNet with more than 200 layers with respect to the training cost. Lastly, one should randomly inspect some feature maps to ensure that all hyperparameter setting works as expected. If a feature map clearly shows no structural correspondence to its input volume and exhibits strong noise or abnormal features, it might signal the existence of overfitting. In this case, a non-zero value of *λ* in the L2 norm regularization should be empirically examined to avoid overfitting.

### Manifold embedding of energy landscape

To prepare for the energy landscape reconstitution, each bootstrapped 3D volume was juxtaposed with its 3D feature map learned by the 3D autoencoder to form an expanded higher-dimensional data point. Each expanded data point is either low-pass filtered at a given resolution (5 Å in default setting) or standardized by the z-score method prior to t-SNE processing:

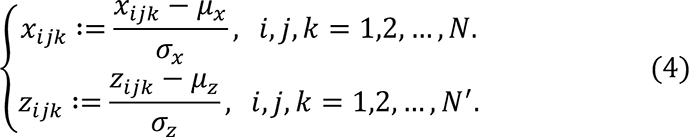

in which *μ_x_* and *σ_x_* denote the mean and standard deviation of the 3D volume voxels *x_ijk_* with the box size of *N*^’^, respectively. Similarly, *μ_z_* and *σ_z_* are the mean and standard deviation of the 3D feature map voxels *z_ijk_* with the box size of *N*^’^, respectively. All the input data points were then embedded onto a low-dimensional manifold via the t-SNE algorithm by preserving the geodesic relationships among all high-dimensional data^51^. During manifold embedding, it is assumed that the pairwise similarities in the high dimensional data space and low dimensional latent space follow a Gaussian distribution and a student’s t-distribution, respectively. To find the low-dimensional latent variable of each data point, the relative entropy, also called the Kullback-Leibler (KL) divergence, is used as an objective function measuring the distance between the similarity distribution *p_ij_*, in the data space and *q_ij_*, in the latent space:

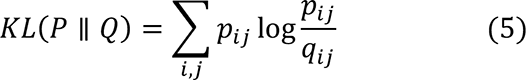

which is minimized by the gradient descent algorithm with the momentum method^69^. The hyperparameter of t-SNE may be tuned to better preserve both local and global geodesic distances (https://distill.pub/2016/misread-tsne/).

The idea of using juxtaposed data composed of both volumes and their feature maps for manifold embedding is similar to the design philosophy of deep residual learning, in which the input and feature output of a residual learning block are added together to be used as input of the next residual learning block. Such a design has been demonstrated to improve learning accuracy and reduce the performance degradation issues when the neural network goes much deeper^49, 50^. Although the 3D feature maps or the bootstrapped volumes alone can be used for manifold embedding, both appear to be inferior in 3D classification accuracy (Supplementary Fig. 7). The juxtaposed format of input data for manifold learning is potentially beneficial to the applications in those challenging scenarios, such as visualizing a reversibly bound small protein like ubiquitin (∼8.6 kDa) of low occupancy by the focused AlphaCryo4D classification (see below).

After the manifold embedding by t-SNR, each 3D volume is mapped to a low-dimensional data point in the learned manifold. The coordinate system, in which the low-dimensional representation of the manifold is embedded, is used for reconstructing energy landscape. The difference in the Gibbs free energy *ΔG* between two states with classified particle numbers of *N_i_* and *N_j_* is defined by the Boltzmann relation *N_i_*/*N_j_* = exp(−*ΔG*/*k_B_T*). Thus, the free energy of each volume can be estimated using its corresponding particle number:

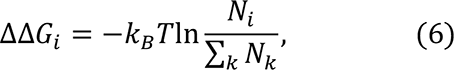

where ΔΔ*G_i_* denotes the free energy difference of the data point with the classified particle number of *N*_i_ against a common reference energy level, *k_B_* is the Boltzmann constant and *T* is the temperature in Kelvin. The energy landscape was plotted by interpolation of the free energy difference in areas with sparse data. We suggest that linear interpolation is used for the energy landscape with loosely sampled areas to avoid overfitting by polynomial or quadratic interpolation. For densely sampled energy landscape, polynomial interpolation could give rise to smoother energy landscape that is easier to be tackled by the string method for MEP solution (see below).

The coordinate system of the embedded manifold output by t-SNE is inherited by the reconstitution of energy landscape as reaction coordinates. They do not have intuitive physical meaning of length scale in the real space. However, they can be viewed as transformed, rescaled, reprojected coordinates from the real-space reaction coordinates along which the most prominent structural changes can be observed. Alternatively, they can be intuitively understood as being similar to transformed, reprojected principal components (PCs) in principle component analysis (PCA). In the case of the 26S proteasome, the two most prominent motions are the rotation of the lid relative to the base and the intrinsic motion of AAA-ATPase motor^23^, which can be approximately projected to two reaction coordinates. In real space, both motions are notably complex and in fact are characterized by a considerable number of degrees of freedom, which are partially defined by the solved conformers.

### String method for finding minimum energy path (MEP)

The string method is an effective algorithm to find the MEP on the potential energy surface^52^. To extract the dynamic information implicated in the experimental energy landscape, an improved and simplified version of the string method has been previously developed^53^. Along the MEP on the energy landscape, the local minima of interest could be defined as 3D clustering centers to guide the particle-voting algorithm for 3D classification to generate high-resolution cryo-EM density maps (Supplementary Fig. 1b). The objective of the MEP identification in energy barrier-crossing events lies in finding a curve *γ* having the same tangent direction as the gradient of energy surface ∇*G*. It can be expressed as:

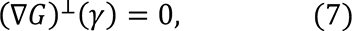

where (∇*G*)⊥ denotes the component of ∇*G* perpendicular to the path *γ*. To optimize this objective function, two computational steps, referred to as ‘evolution of the transition path’ and ‘reparameterization of the string’, are iterated until convergence within a given precision threshold.

After initialization with the starting and ending points, in the step of evolution of the transition path, the positions of interval points along the transition path were updated according to gradient of the free energy at the *t*th iteration:

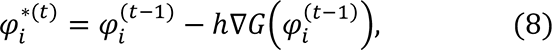

with 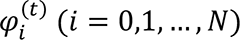 being the *i*th intermediate point along the transition path at the *t*th iteration (∗ denoting the temporary values), and ℎ the learning rate.

In the step of reparameterization of the string, the values of positions 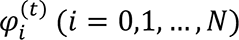 were interpolated onto a uniform mesh with the constant number of points. Prior to interpolation, the normalized length 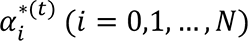 along the path was calculated as:

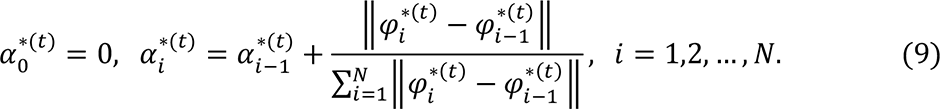

Given a set of data points 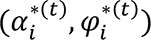, the linear interpolation function was next used to generate the new values of positions 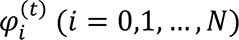at the uniform grid points 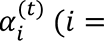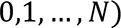. The iteration was terminated when the relative difference 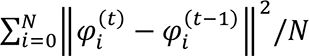 became small enough.

### String method parameters

The string method of searching a rational MEP on the energy landscape can only guarantee the solution of local optimum and is not capable of ensuring a solution of global optimum^53^. The outcome of the string method depends on the initialization of the starting and ending points of the MEP on the energy landscape. If there are too many local minima to sample along a long MEP, the string method could retrieve a MEP solution that partly misses some local energy minima by going off the pathway. In this case, the search of an expected MEP solution can be divided into several segments of shorter MEPs connecting one another, with each MEP being defined by a closer pair of starting and ending points, which travels through a smaller number of energy minima. Another parameter affecting the MEP solution is the step size being set to explore the MEP on energy landscapes. Although a default value of 0.1 empirically recommended may work in many cases, it may be helpful to tune the step size according to the quality of the energy landscape. The value of step size can be decreased if the computed path runs out of the energy landscape and can be increased if the path updates too slowly during iterative searching by the string method.

### Energy-based particle voting algorithm

The particle-voting algorithm was designed to conduct 3D classification, particle quality control, reproducibility test and particle selection in an integrative manner. The particle-voting algorithm mainly involves two steps (Supplementary Fig. 1b). First, we count the number of votes for each particle mapped within the voting boundaries of all 3D clusters on the energy landscape. One vote is rigorously mapped to one copy of the particle used in reconstructing a 3D volume and to no more than one 3D cluster on the energy landscape where the corresponding volume is located. Thus, each particle can have *M* votes casted for no more than *M* 3D clusters. If the vote is mapped outside of any 3D cluster boundary, it becomes an ‘empty vote’ with no cluster label. Each non-empty vote is thus labeled for both its particle identify and corresponding cluster identity. For each pair of particle and cluster, we compute the total number (*V*) of votes that the cluster receives from the same particle. Each particle is then assigned and classified to the 3D cluster that receives *V* > *M*/2 votes from this particle (Supplementary Fig. 1b). Note that after particle voting, each particle is assigned no more than once to a 3D class, with its redundant particle copies removed from this class. This strategy only retains the particles that can reproducibly vote for a 3D cluster corresponding to a homogeneous conformation, while abandoning those non-reproducible particles with divergent, inconsistent votes.

### Alternative strategy of 3D clustering without particle voting

Because the particle-voting algorithm imposes strong constraints on the reproducibility of particle classification by deep manifold learning, some 3D classes might be assigned with a small number of particles that are insufficient to support high-resolution reconstruction. To remedy this limitation, an alternative, distance-based classification algorithm was devised to replace the particle-voting algorithm when there are not enough particles to gain the advantage of particle voting (Supplementary Fig. 1c). In this method, the distances of all *M* copies of each particle to all 3D cluster centers on the energy landscape are measured and ranked. Then, the particle is classified to the 3D cluster of the shortest distance. A threshold could also be manually preset to remove particles that are too far away from any of the cluster centers. The distance-based classification method can keep more particles, but it ignores the potential issue of irreproducibility of low-quality particles. Thus, it is proven to be less accurate in 3D classification (Supplementary Fig. 7m-o). In other words, it trades off the classification accuracy and class homogeneity to gain more particles, which is expected to be potentially useful for small datasets or small classes. By contrast, the energy-based particle-voting algorithm imposes a more stringent constraint to select particles of high reproducibility during classification, resulting in higher quality and homogeneity in the classified particles, which is superior to the distance-based classification method (Supplementary Fig. 7s-u).

### Practical consideration of particle shuffling and voting parameters

The parameter *M* determines the degree of implicit cross-validation of classification reproducibility, as well as the sampling densities of energy landscape. To establish reproducibility for 3D classification, *M* should be no less than 3. The variation of *M* is not supposed to change the energy landscape because it multiplies on both denominator and enumerator in the Boltzmann relation and is cancelled in the ratio of particle densities in computing the free-energy differences. Increasing *M* value will allow more volumes to be bootstrapped, which then leads to a higher sampling density in computing the manifold of energy landscape and potentially enhances its reconstitution quality. The default classification threshold *M*/2 is the minimal number of votes for verifying the reproducibility of 3D classification. However, a higher threshold, such as 2*M*/3, will give rise to more stringent criteria in cross-validation, with a tradeoff of voting out more particles. The particle voting algorithm does not entirely eradicate misclassified particles. However, increasing either *M* or the classification threshold could theoretically have similar impact and improve the conformational homogeneity, because it gets less probable for a particle being misclassified to the same cluster for more times.

Several considerations may be applied to the choice of *M* and to set up particle shuffling and voting. First, to obtain a high-quality energy landscape, a thousand or more bootstrapped volumes are expected for datasets of more than 1-2 million particles. Second, the average particle number per volume is expected to be no less than 5000 or more to ensure that majority of volumes include sufficient particles for quality reconstructions. The expected particle number per volume must increase if the average SNR per particle is decreased. It may need to reach 10000-20000 or more for small proteins or lower SNR datasets. Third, for a dataset of a moderate size, *M* can be adjusted to a higher value to mitigate the lack of image data for bootstrap.

### Data-processing workflow of AlphaCryo4D

The following summarizes the complete procedure of AlphaCryo4D, with the detailed algorithm rationale explained in the previous subsections.

**Input**: Single-particle cryo-EM dataset after initial particle rejection of apparent false particles.

**Output**: Energy landscape, MEP, 3D class assignment of each particle.

**Step 1**. Bootstrap many 3D volumes through particle shuffling, consensus alignment and Bayesian clustering.

1. Split the particle dataset randomly to many sub-datasets, if necessary, in case of a large dataset (e.g., >150,000 particle images), for batch processing of particle shuffling and volume bootstrapping. Otherwise, skip this step if the dataset is small enough (e.g., <150,000).
2. For each sub-dataset, conduct a consensus alignment to generate initial parameters of Euler angles and translations in RELION with one or more consensus models.
3. Divide each sub-dataset into M + 1 groups, shuffle the sub-dataset M + 1 times and each time take a different group out of the shuffled sub-dataset, giving rise to M + 1 shuffled sub-datasets all with different collection of particles.
4. Conduct 3D Bayesian classification on all the M + 1 shuffled sub-datasets to generate hundreds of 3D volumes, making each particle to contribute to M different volumes.
5. Execute steps (2) to (4) using the same initial model (low-pass filtered at 60-Å) for all sub-datasets.

**Step 2**. Extract deep features of all volume data with the 3D deep residual autoencoder.

6. Align all 3D volumes and adjust them to share a common frame of reference.
7. Initialize the hyper-parameters of the 3D autoencoder (Supplementary Table 1).
8. Train the neural network with the 3D volume data to minimize mean square error between the decoding layer and the input by the Adam algorithm of initial learning rate 0.01.
9. Extract the 3D feature maps of all volumes from the encoding layer.

**Step 3**. Embed the volume data to two-dimensional manifolds through the t-SNE algorithm and compute the energy landscape and find the MEP.

10. Calculate the pairwise similarities between volumes using their feature-map-expanded volume vectors, and randomly initialize the low-dimensional points.
11. Minimize the Kullback-Leibler divergence by the Momentum algorithm to generate 2D manifold embeddings with t-SNE.
12. Compute the energy landscape from the manifold using the Boltzmann relation.
13. Initialize searching of the MEP with a straight line between given starting and ending points.
14. Find the optimal MEP solution using the string method.

**Step 4**. Classify all particles through the energy-based particle-voting algorithm.

15. Sample the clustering centers along the MEP and calculate the recommended clustering radius.
16. Define the local energy minima as the 3D clustering centers and their corresponding cluster boundary for particle voting.
17. For each particle, cast a labelled vote for a 3D class when a volume containing one of the *M* particle copies is located within the voting boundary.
18. Count the number of votes of each particle with respect to each 3D class and assign the particle to the 3D class that receives more than *M*/2 votes from this particle.
19. Refine each 3D density map separately to high resolution in RELION or cryoSPARC using particles classified into the same 3D classes.

### Blind assessments with simulated datasets

Three simulated large datasets with the SNRs of 0.05, 0.01 and 0.005, each including 2-million particle images, were employed to benchmark AlphaCryo4D and to compare its performance with alternative methods. For each synthetic dataset, the particles were computationally simulated by projecting the 20 3D density maps calculated from 20 hypothetical atomic models emulating continuous rotation of the NLRP3 inflammasome protein. The 20 atomic models were interpolated between the inactive NLRP3 structure and its hypothetical active state, which was generated through homology modeling using the activated NLRC4 structure^54^ (Fig. 1c). The 20 atomic models represent sequential intermediate conformations during a continuous rotation in its NATCH domain against its LRR domain over an angular range of 90°. Each conformation is thus rotated 4.5° over its immediate predecessor in the conformational continuum sequence. 100,000 simulated particle images per conformational state were generated with random defocus values in the range of −0.5 to −3.0 μm, resulting in 2 million particles for each dataset of a given SNR. The pixel size of the simulated image was set to the same as the pixel size (0.84 Å) of the real experimental dataset of NLRP3-NEK7 complex^54^. To emulate realistic circumstances in cryo-EM imaging, Gaussian noises, random Euler angles covering half a sphere and random in-plane translational shifts from −5.0 to 5.0 pixels were then applied to every particle image.

Each simulated heterogeneous NLRP3 dataset was analyzed separately by AlphaCryo4D and used to characterize the performance and robustness of AlphaCryo4D against the variation of SNRs. In the step of particle shuffling and volume bootstrapping, 2,000,000 particles in the dataset of any given SNR were divided randomly into 10 sub-datasets for batch processing. The orientation of each particle was determined in the initial 3D consensus alignment in RELION, which did not change in the subsequent 3D classification. In this step, the maximum number of iterations of the 3D alignment was set up as 30, with the initial reference low-pass filtered to 60 Å. 3-fold particle shuffling (indicated as × 3 below) was conducted on each sub-dataset for volume bootstrapping. The first round of maximum-likelihood 3D classification divided the input shuffled particle sub-dataset into 5 classes, each of these classes were then further classified into 8 classes. This procedure was separately executed on all shuffled particle sub-datasets. The particle shuffling and volume bootstrapping generated 1,372, 1,489, and 1,587 volumes by the datasets with SNRs of 0.05, 0.01 and 0.005, respectively. These volume data were used as inputs for deep residual autoencoder to compute low-dimensional manifolds and energy landscapes (Fig. 1d-f, Supplementary Fig. 1b, 2a, b). After searching the MEP on the energy landscapes by the string method, 20 cluster centers along the MEP were defined by the local energy minima along the MEP by approximately equal geodesic distance between adjacent minima, which represent potentially different conformations of the molecule (Supplementary Fig. 2a-c). The particle-voting algorithm was applied in every cluster to determine the final particle sets for all 3D classes. For validation of the methodology and investigation of 3D classification improvement, we labeled each bootstrapped 3D volume with the conformational state that held the maximum proportion of particles in the class and computed its 3D classification precision as the ratio of the particle number belonging to the labelled class versus the total particle number in the volume (Fig. 2a, Supplementary Fig. 3a, b).

### Analysis of algorithm mechanism

To understand how the 3D classification accuracy is improved by AlphaCryo4D, we analyzed the algorithmic mechanism by a number of control experiments (Supplementary Fig. 2d-g, 7). In total, we conducted 24 conditional tests using 10 variations of algorithmic design by removing or replacing certain components of AlphaCryo4D. First, by removing the entire component of deep manifold learning, the distributions of high-precision 3D classes obtained by *M*-fold particle shuffling and voting alone drop by ∼60% relative to that from complete AlphaCryo4D processing (Supplementary Fig. 7d-f). In these control experiments, the particle voting was achieved through counting the votes of a particle against the same conformers classified by RELION via computing the Intersection-over-Union (IoU) metric after the step of volume bootstrapping. Similarly, by removing the 3D deep residual autoencoder but keeping the manifold learning by t-SNE for energy landscape reconstitution, the distributions of high-precision 3D classes is reduced by ∼15% at the SNR of 0.005 (Supplementary Fig. 7p-r). The effect of accuracy degradation is less prominent at a higher SNR (0.05 or 0.01), indicating that the unsupervised feature learning using deep residual autoencoder promotes the tolerance of the algorithm against higher noise level in the data. Together, these control tests suggest that deep manifold learning plays a crucial role in improving 3D classification accuracy with low SNR data.

Next, we replaced t-SNE with four other algorithms in manifold learning step, including two classic linear dimensionality reduction techniques, principal component analysis (PCA) ^13^ and multidimensional scaling (MDS) ^55^, and two nonlinear dimensionality reduction algorithms, isometric mapping (Isomap) ^56^ and locally linear embedding (LLE) ^57^. We applied the four algorithms to reduce the dimensionality of the same sets of bootstrapped volume data. Although being capable of differentiating a portion of ground-truth conformers, the resulting 2D mappings by the four techniques exhibit considerable overlap between adjacent conformers and are incapable of distinguishing all 20 conformers of ground truth, thus inevitably missing many conformers (Supplementary Fig. 2d-g). The PCA and Isomap approximately missed half of the ground-truth conformers, whereas MDS and LLE both missed around 70% of the ground-truth conformers. The inferior performance of these techniques is consistent with the original control experiments conducted by the t-SNE developers ^51^.

Further, we conducted control experiments to evaluate the impact of particle voting on the 3D classification accuracy by replacing the particle voting component with a clustering strategy that directly classifies each particle to the cluster of the nearest clustering center (Supplementary Fig. 1c). In this case, the distributions of high-precision 3D classes are reduced by ∼15-30% (Supplementary Fig. 7m-o). The reduction is more prominent at the lower SNR. This indicates that the particle voting considerably improves the 3D classification accuracy but is less impactful than the component of deep manifold learning.

Last, by tracking the statistics of classification precisions step by step, we evaluated how the 3D classification accuracy is improved over the intermediate steps of AlphaCryo4D (Supplementary Fig. 7g-l). We found that the steps of particle shuffling, defining cluster boundaries on the energy landscapes and energy-based particle voting contribute to ∼16%, ∼20% and ∼40% improvements in the distributions of high-precision 3D classes, respectively. Taken together, these results indicate that all components contribute to the improved performance of AlphaCryo4D in 3D classification toward the atomic level. Neither deep manifold learning nor particle voting alone is sufficient to achieve the present level of 3D classification accuracy.

### Comparisons with alternative methods

Using 3D classification precision as a benchmark indicator, the performance of AlphaCryo4D preprocessed by standardization or 5 Å low-pass filtering was compared with several other methods: (1) the conventional maximum-likelihood-based 3D (ML3D) classification in RELION^3, 4, 6^ with and without a hierarchical strategy, (2) the conventional heterogeneous reconstruction in cryoSPARC^2^, (3) the 3DVA algorithm with 2 and 3 principal components (PCs) in cryoSPARC^11^ and (4) the deep generative model-based cryoDRGN^12^. A total of 24 comparative tests by these alternative methods have been conducted blindly using the three simulated datasets, which includes 6-million images in total. In all our comparative tests on AlphaCryo4D and the alternative methods, the ground-truth information of particle orientations, in-plane translations and conformational identities were completely removed from the methods being tested and were not used for any steps of data processing or training. The ground-truth conformational identities of particles were only used as validation references to compute the 3D classification precisions of the blind testing results.

For the tests using ML3D in RELION, we classified all particles directly into 20 classes and hierarchically into 4 × 5 classes, which first divided the dataset into 4 classes, with each class further classified into five sub-classes (Supplementary Fig. 3g, h). For testing the conventional discrete ab initio heterogeneous reconstructions in cryoSPARC, each synthetic dataset was directly classified into 20 classes without providing any low-resolution reference model (Supplementary Fig. 3f). For comparison with the 3DVA algorithm in cryoSPARC, we first did the blind consensus alignment of the entire dataset to find the orientation of each particle. Then the alignment and the mask generated from the consensus reconstruction were used as inputs into the 3DVA calculation, with the number of orthogonal principal modes being set to 2 or 3 in 3D classification. The 3D variability display module in the cluster mode was used to analyze the results of 3D classification. For blind tests with cryoDRGN, a default 8D latent variable model was trained for 25 epochs. The encoder and decoder architectures were 256 × 3, as recommend by the cryoDRGN developers^12^. The particle alignment parameters prior to cryoDRGN training were obtained by the same blind consensus refinement in RELION used for other parallel tests. The metadata of 3D classification precisions as well as the 3D density maps from all the algorithms applied on the three simulated datasets were collected to conduct the statistical analysis (Fig. 2 and Supplementary Fig. 3).

### Computational costs

Although the computational cost of AlpahCryo4D is generally higher than the conventional approach, it does not appear to increase drastically and likely falls in an affordable range, while reducing the average cost of computation per conformational state. In a nutshell, we can have a brief comparison of the computational efficiency on the simulated 2-million image dataset with an SNR of 0.01. In the step of 3D data bootstrapping, we split the dataset into 20 subsets, which contained 100,000 particles each. The 3D consensus alignment of all 2,000,000 particles cost about 75 hours using 8 Tesla V100 GPUs interconnected with the high-speed NVLink data bridge in a NVIDIA DGX-1 supercomputing system. Within each subset, the 3D Bayesian classification for one leave-one-group-out dataset cost about 2.5 hours using 320 CPU cores (Intel Xeon Gold 6142, 2.6 GHz, 16-core chip), so the total time spent in one subset was about 10 hours using 320 CPU cores in an Intel processor-based HPC cluster. In addition, it spent about 3 hours to extract feature via deep neural network using 8 Tesla V100 GPUs of the NVIDIA DGX-1 system. In contrast to about 160 hours cost in traditional classification methods, this approach cost a little more than 200 hours using 8 Tesla V100 GPUs and 320 CPU cores. Taken together, these observations suggest that the computational cost and efficiency of AlphaCryo4D are within the acceptable range considering the output of more high-resolution conformers yielded by the procedures. The benefits likely outweigh the tradeoff of computational costs, particularly for large datasets that equally slow down any methods.

### Applications to the experimental cryo-EM dataset of the membrane channel protein complex RyR1

To determine the dynamics of the peripheral domains of the RyR1, density subtraction and focused energy landscape were applied in the further analysis of state 1 conformation^45^. Due to the low SNR of the particles, a large *M* = 30 was used in the particle shuffling in AlphaCryo4D. After symmetry expansion of particles of state 1, a total of 1,557,168 particles with the core domain subtracted were utilized to compute the energy landscape with 310 volumes. Three 3D classes with the particle numbers of 196,669, 848,655 and 189,796 were obtained by the particle-voting procedure on the energy landscape. All these 3 classes were merged together for high resolution refinement with a mask of the peripheral domains using RELION. To further improve the quality of the density map, a zoom-in energy landscape with *M* = 22 was computed to result in a final particle set with the number of 1,198,946. Furthermore, as the P2 domain showed high flexibility, we subtracted the non-P2 high-resolution components from the particles and computed the P2 domain-focused energy landscape using *M* = 23. This energy landscape resulted in three classes with the particle numbers of 14,089, 1,108,282 and 57,381, which was combined to refine the final density map of the P2 domain with a specific mask using RELION.

### Applications to the experimental cryo-EM dataset of the human 26S proteasome

The substrate-engaged human 26S proteasome dataset^23^ (EMPIAR-10669) includes 3,254,352 RP-CP particles (combined with particle images from both the DC and SC proteasomes) in total, with the super-resolution counting mode pixel size of 0.685 Å and the undecimated box size of 600 × 600 pixels. The substrates of 26S proteasome first appear in the state E_A2_. In this focused study, 147,108 particles of E_A2_ were utilized for computing the specific energy landscape with 40 volumes. The alignment of these particles was firstly refined with a 19S mask. *M* = 3 particle shuffling was then conducted to bootstrap the 3D volumes with a soft mask of 19S. By clustering on the energy landscape and particle voting, two classes containing 99,043 and 47,389 particles were generated for high resolution refinement using RELION, which were designated state E_A2.1_ and E_A2.0_, respectively. The density map of state E_A2.0_ exhibits a conformation identical to state E_A2_ in the previous report^23^.

### Applications to the experimental cryo-EM dataset of the *Pf*80S ribosome

The *Pf*80S ribosome dataset (EMPIAR-10028) contains 105,247 particle images. First, the alignment of all particles was refined with a global mask in RELION. To bootstrap enough 3D volumes for the energy landscape, we set *M* = 7 in the particle shuffling step, resulting in 79 volumes in total with a global mask and 15 Å resolution limit in the expectation step. Moreover, the regularization coefficient was set to 0.001 when training the 3D residual network with this small dataset. All volumes were then low-pass filtered at 5 Å prior to manifold embedding. Five clusters were obtained from the energy landscape, which had 66,035, 21,482, 9,922, 6,424, 657 particle images respectively after voting. All these clusters were refined independently using RELION.

### Applications to the experimental cryo-EM dataset of the pre-catalytic spliceosome

The pre-catalytic spliceosome dataset (EMPIAR-10180) with the particle number of 327,490 shows high conformational dynamics. The consensus alignment of these particles in the original dataset was used in the first step. *M* = 7 particle shuffling was utilized to bootstrap 160 volumes with a soft global mask and 15 Å resolution limit in the expectation step. The regularization coefficient of 0.001 was set to train the 3D Autoencoder with this dataset. All volumes were then low-pass filtered at 5 Å prior to manifold embedding. Based on the energy landscape, we obtained 14 classes with the particle number of 15,749, 17,703, 27,522, 13,820, 20,841, 14,989, 16,377, 22,047, 25,049, 21,327, 18,949, 6,157, 9,790 and 36,878 after voting. Each class was refined independently using RELION.

### Applications to the experimental cryo-EM dataset of the bacterial 50S ribosomal intermediates

The 131,899 particle images of the 50S ribosomal large subunit (EMPIAR-10076) are highly heterogeneous. We refined the alignment of all particles with a global mask of 50S using RELION. In the first step, 119 bootstrapped volumes were utilized to calculate the energy landscape. *M* = 7 particle shuffling was conducted to generate these volumes without any mask. The regularization coefficient of 0.001 was set when training the deep neural network. All volumes were then low-pass filtered at 5 Å prior to manifold embedding. Then the first energy landscape resulted in nine clusters with the particle number of 766 (A), 15,129 (B), 1,236, 322, 670, 22,037, 25,445 (D), 33,976 (E) and 25,115 (C) respectively. For the clusters of states B-E, four zoom-in energy landscapes were then computed to discover more sub-states with 46, 56, 66 and 72 bootstrapped volumes respectively. In the zoom-in energy landscape calculation, *M* = 7 was set in particle shuffling for bootstrapping 3D volumes with a global mask and 20 Å resolution limit in the expectation step. Then the regularization coefficient of 0.001 was used to train the deep residual network. All volumes were then low-pass filtered at 5 Å prior to manifold embedding. After particle voting, the energy landscape of state B resulted in three sub-clusters with the particle number of 3,494, 3,600 and 7,803. The state C energy landscape resulted in five sub-clusters with the particle number of 2,444, 7,442, 711, 7,633 and 5,557. The state D energy landscape resulted in five sub-clusters with the particle number of 6,231, 8,785, 2,123, 3,718 and 4,092. And the energy landscape of state E resulted in eight sub-clusters with the particle number of 4,783, 1,588, 5,891, 3,620, 7,352, 5,265, 357 and 2,241. All these clusters were refined using RELION.

## Code availability

Source code of AlphaCryo4D is freely available at http://github.com/alphacryo4d/alphacryo4d/.

## Acknowledgments

The authors thank Q. Ouyang and W. L. Wang for constructive discussions. This work was funded in part by the Beijing Natural Science Foundation (grant No. Z180016/Z18J008), the National Natural Science Foundation of China (grant No. 11774012) and a gift academic grant from Intel Corporation.

## Author contributions

Y.Mao and Z.W. conceived this study and devised the methodology. Z.W. developed the prototype source code of the AlphaCryo4D system, conducted numerical studies of the system using the synthetic datasets. Z.W., E.C., and S.Z. analyzed the experimental cryo-EM datasets and refined the density maps. C.L. and C.C.Yin contributed to the sample preparation and data collection on the RyR1 dataset. Y.Ma contributed to early studies in algorithm design and testing. Y.Mao supervised this study, analyzed the data and wrote the manuscript with inputs from all authors.

## Additional information

**Supplementary information** is available for this paper online.

**Correspondence and requests for materials** should be addressed to Y.M.

---

**Supplementary Fig. 1.**
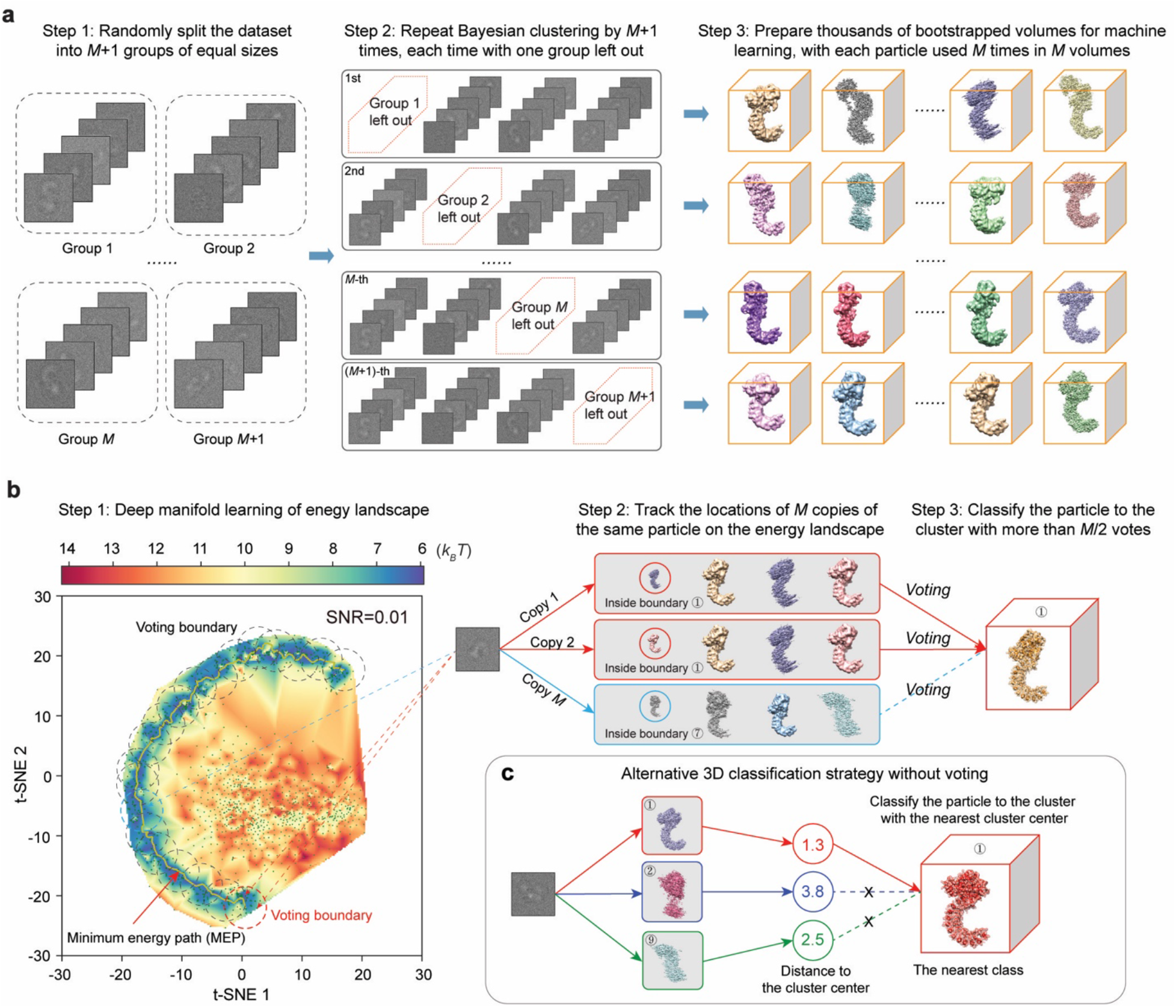
Detailed algorithmic design of particle shuffling and voting in AlphaCryo4D. **a,** Schematic showing the method of *M*-fold particle shuffling and bootstrapping of 3D volumes. In the step 1, all particles are split randomly into *M* + 1 groups equally. Then the step 2 carries out the Bayesian clustering for reconstructions of 3D density maps within each of the *M* + 1 particle sets that are shuffled via the leave-one-group-out approach. After these two steps, thousands of 3D volumes are generated for the subsequent 3D deep learning, with each particle contributing to *M* volumes. **b,** Schematic showing the algorithmic concept of energy-based particle voting for 3D classification. The left panel shows the energy landscape obtained by deep manifold learning. After clustering along the minimum energy path, all *M* locations of each particle on the energy landscape can be tracked to cast *M* votes. A vote of the particle is only counted for the cluster when it is mapped within the voting boundary of the cluster, as indicated by the circles marked on the energy landscape. Eventually, this particle is classified to the 3D cluster with over *M*/2 votes from this particle. **c,** Alternative distance-based 3D classification method as a control in the analysis of algorithmic performance of particle voting. Instead of particle voting, each particle is directly classified to the cluster with the nearest clustering center among the M volume data points in the distance-based 3D classification strategy.

**Supplementary Fig. 2.**
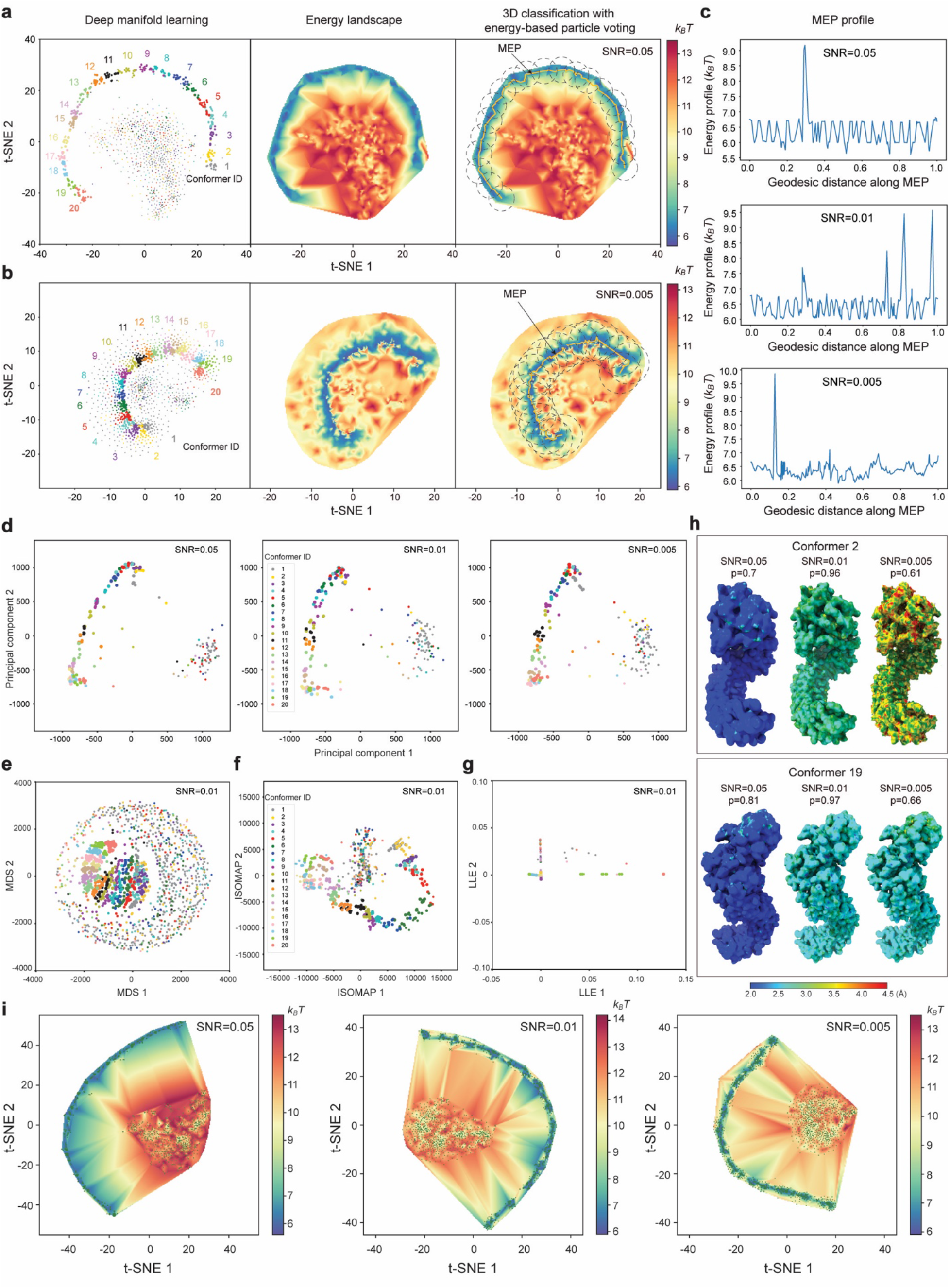
Blind assessments of AlphaCryo4D and its comparison with the 3D PCA method using the simulated heterogeneous NLRP3 datasets of different SNRs. **a** and **b,** Reconstruction of the energy landscape of the simulated NLRP3 datasets at SNRs of 0.05 (**a**) and 0.005 (**b**) by the t-SNE algorithm using the bootstrapped volumes and their corresponding feature maps. Colors in the left panels indicate the ground truth of 3D volume data points. **c,** Free energy profiles along the MEP calculated by the string method in the 2D energy landscapes of the simulated NLRP3 datasets at SNRs of 0.05 (top), 0.01 (middle) and 0.005 (bottom). **d,** Linear dimensionality reduction of bootstrapped 3D volumes from the simulated datasets at three distinct SNRs by the 3D PCA method. **e**-**g,** Dimensionality reduction of bootstrapped 3D volumes from the simulated dataset at SNR of 0.01 by the multidimensional scaling (MDS)^55^ (panel **e**), isometric mapping (Isomap)^56^ (panel **f**) and locally linear embedding (LLE)^57^ algorithms (panel **g**). Colors of data points indicate the ground truth of their corresponding 3D volumes. **h,** Comparison of local resolution assessment of AlphaCryo4D-classified NLRP3 reconstructions of conformers 2 (upper inset) and 19 (lower inset) from the simulated datasets of three distinct SNRs. The local resolutions were computed by ResMap^70^. Conformers 2 and 19 have been missed 10 and 8 times in 18 control tests using several other methods (see Supplementary Fig. 3c-h), which are the most and second-most frequently missed conformers, respectively. The 3D classification precision (P) is labelled above each density map. The color bar of local resolution is shown in the lower insert. **i,** Energy landscape calculated by 3D volumes with preprocessing of 5 Å low-pass filtering instead of standardization. The gap between the majority and minority data points was widened at different SNRs.

**Supplementary Fig. 3.**
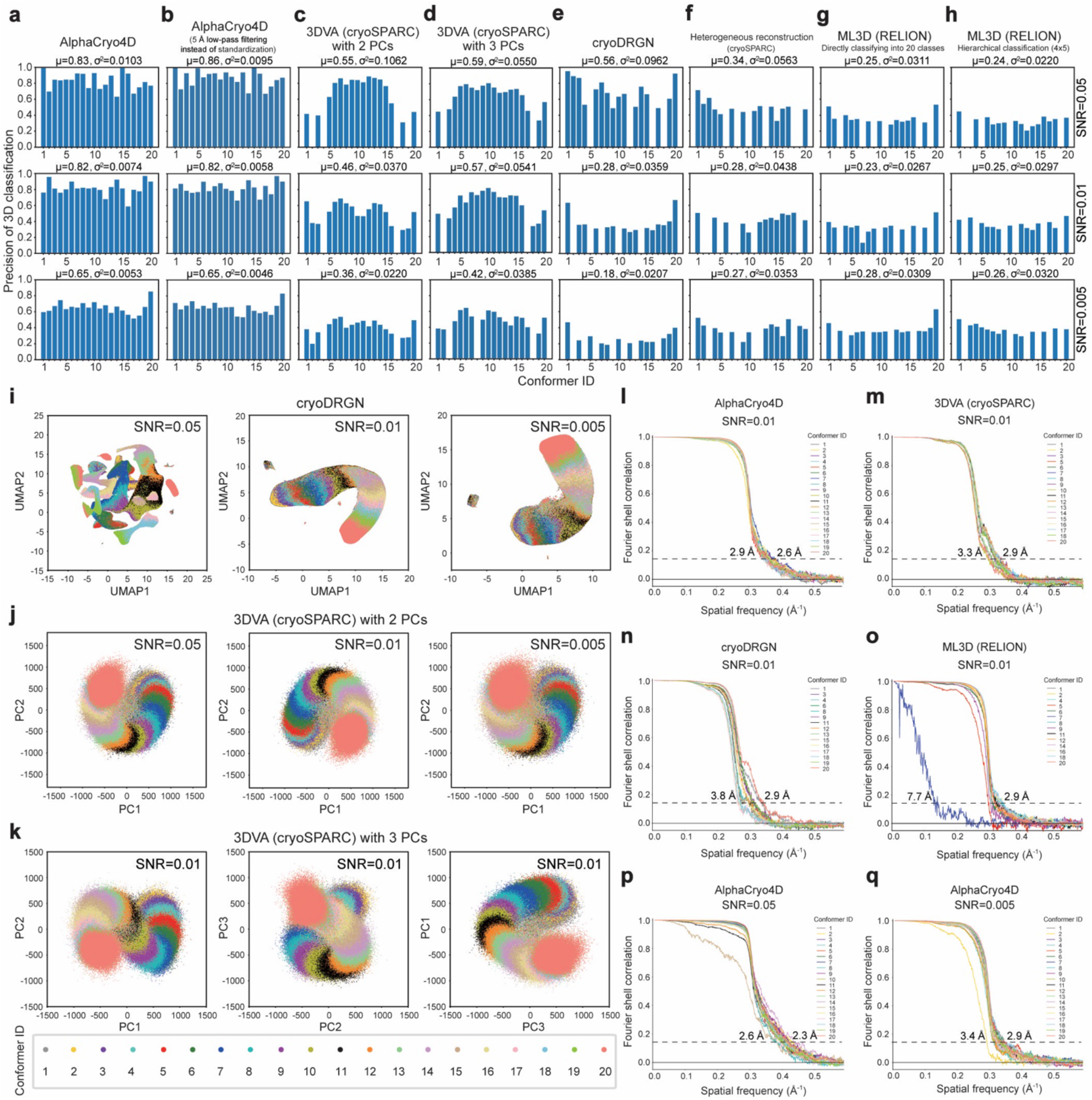
Performance comparison of AlphaCryo4D with alternative methods using the simulated heterogeneous NLRP3 datasets of different SNRs. **a-h,** 3D classification precision of the simulated datasets by AlphaCryo4D (**a**), AlphaCryo4D preprocessed by 5 Å low-pass filtering instead of standardization (**b**), 3DVA with two principal components (PCs) in cryoSPARC^11^ (**c**), 3DVA with three principal components (PCs) in cryoSPARC^11^ (**d**), cryoDRGN^12^ (**e**), heterogeneous reconstruction in cryoSPARC (**f**) and maximum-likelihood classification in RELION (**g**, **h**). The results of SNRs of 0.05 (the first row), 0.01 (the second row) and 0.005 (the third row) are shown on three rows for side-by-side comparison. On the top of each panel, the symbols of µ and *σ*^2^ denote the mean and variance of precision, respectively, with the values of missing classes treated as zeros. In the maximum-likelihood classification of RELION, both direct and hierarchical strategies are compared in the study. **i**-**k,** Visualization of 3D classification by cryoDRGN in autoencoder-learned feature space (**i**), and by 3DVA with 2 PCs (**j**) and 3 PCs (**k**). **l**-**o,** The gold-standard FSC plots of the 20 maps resulting from 3D classification by AlphaCryo4D (**l**), of the 18 maps resulting from the 3D classification by 3DVA in cryoSPARC (**m**), of the 15 maps resulting from cryoDRGN (**n**) and of the 14 maps resulting from the maximum-likelihood 3D classification in RELION (**o**) on the simulated data of 0.01 SNR. They correspond to the precision results presented in the second row of panels (**a**), (**c**), (**e**) and (**g**), respectively. **p** and **q,** Additional gold-standard FSC plots of the refined density maps resulting from AlphaCryo4D on the simulated datasets of SNRs of 0.05 and 0.005.

**Supplementary Fig. 4.**
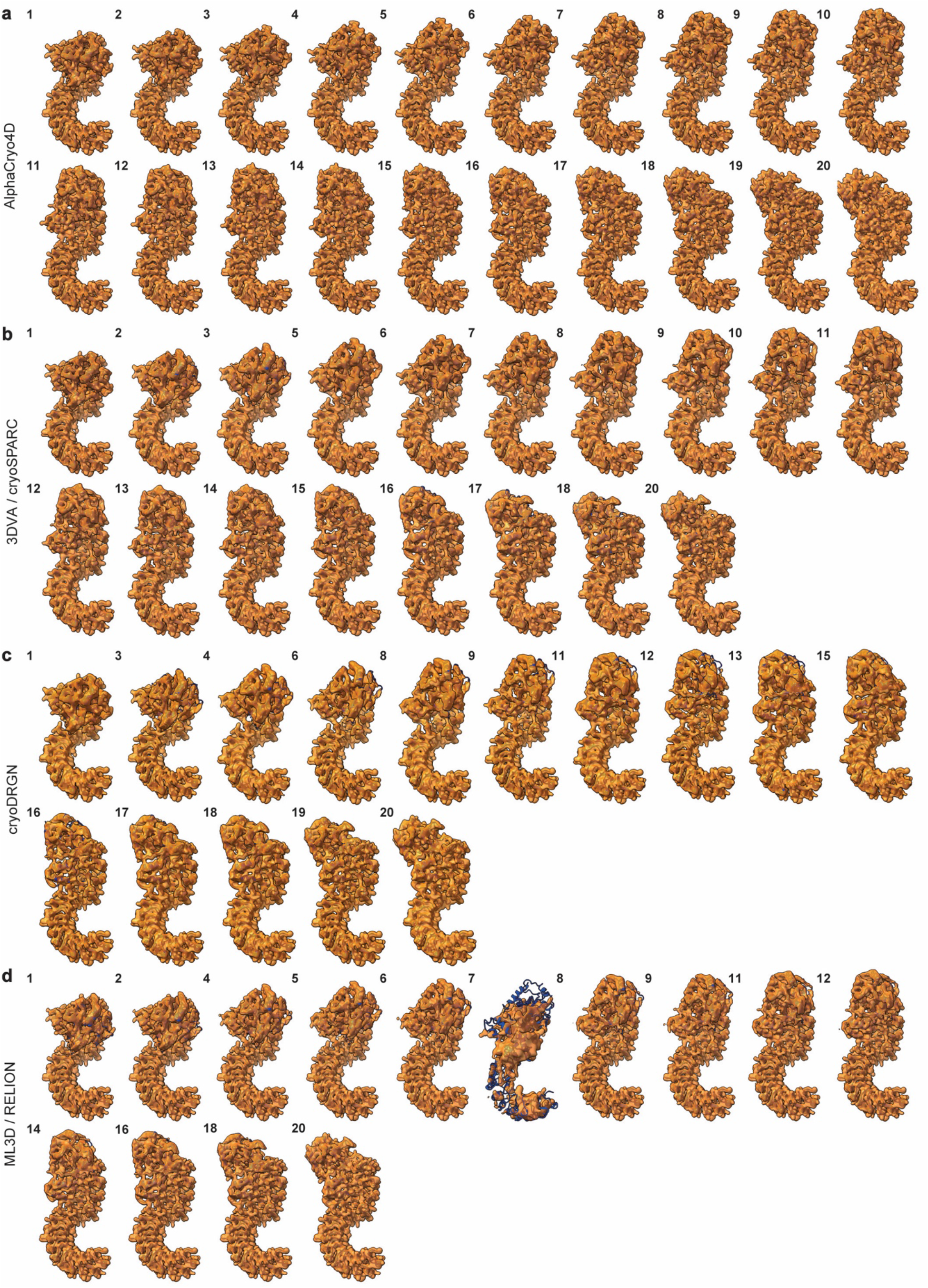
Map assessments of AlphaCryo4D in reconstructions of conformational continuum in comparison with conventional methods on the simulated data of 0.01 SNR. **a,** The 20 maps of distinct NLRP3 conformers resulting from the 3D classification by AlphaCryo4D. **b,** The 18 maps of NLRP3 resulting from 3DVA in cryoSPARC. **c,** The 15 maps of NLRP3 resulting from the 3D classification by cryoDRGN. **d,** The 14 maps of NLRP3 resulting from the 3D classification by ML3D in RELION. All maps are shown in transparent surface representations superimposed with their corresponding atomic models of the ground truth in cartoon representations, which are fitted to the maps as rigid bodies without further atomic modelling. The conformer ID numbers are marked on the upper left of each map panel. The results shown in panels (**a**), (**b**), (**c**) and (**d**) correspond to the FSC results shown in panels (**l**), (**m**), (**n**) and (**o**) of Supplementary Fig. 3, respectively.

**Supplementary Fig. 5.**
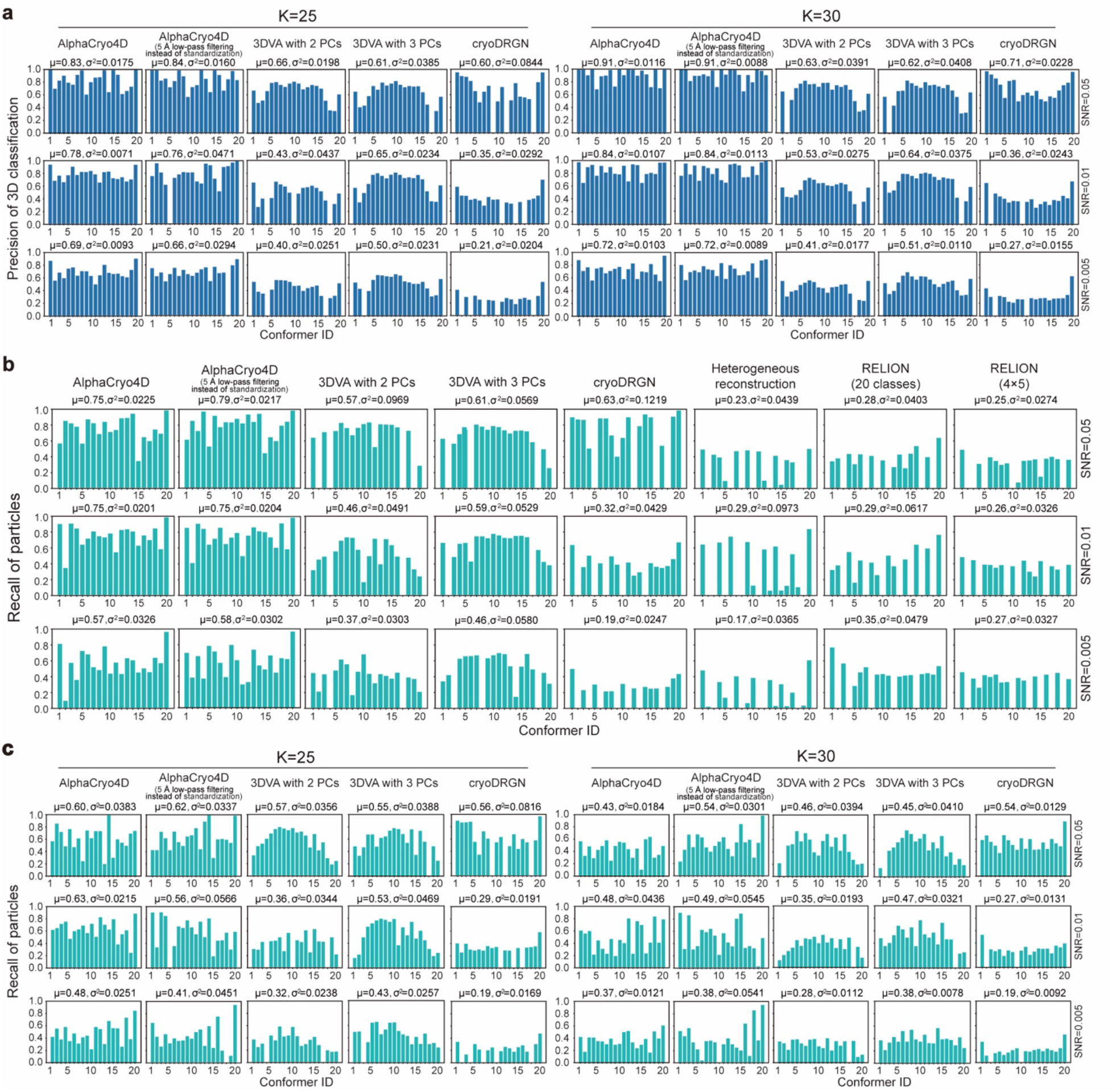
Performance evaluation of 3D classification with alternative parameters and comparison of the recall of 3D classification using the simulated datasets of three distinct SNRs. **a,** 3D classification precision of AlphaCryo4D preprocessed by standardization or 5 Å low-pass filtering, 3DVA using two PCs, 3DVA using three PCs and cryoDRGN when the class number K was set to 25 or 30, which are more than the class number 20 of ground truths. **b,** 3D classification recall of AlphaCryo4D preprocessed by standardization or 5 Å low-pass filtering, 3DVA using two PCs, 3DVA using three PCs, cryoDRGN, heterogeneous reconstruction of cryoSPARC and maximum-likelihood classification of RELION at SNRs of 0.05 (the first row), 0.01 (the second row) and 0.005 (the third row). Particles recall of each state was calculated as the ratio of true positive particles in all ground truth particles. In all methods, the class number was set to 20. **c,** 3D classification recall of AlphaCryo4D preprocessed by standardization or 5 Å low-pass filtering, 3DVA using two PCs, 3DVA using three PCs and cryoDRGN when the class number was set to 25 or 30 (more than the class number 20 of ground truths).

**Supplementary Fig. 6.**
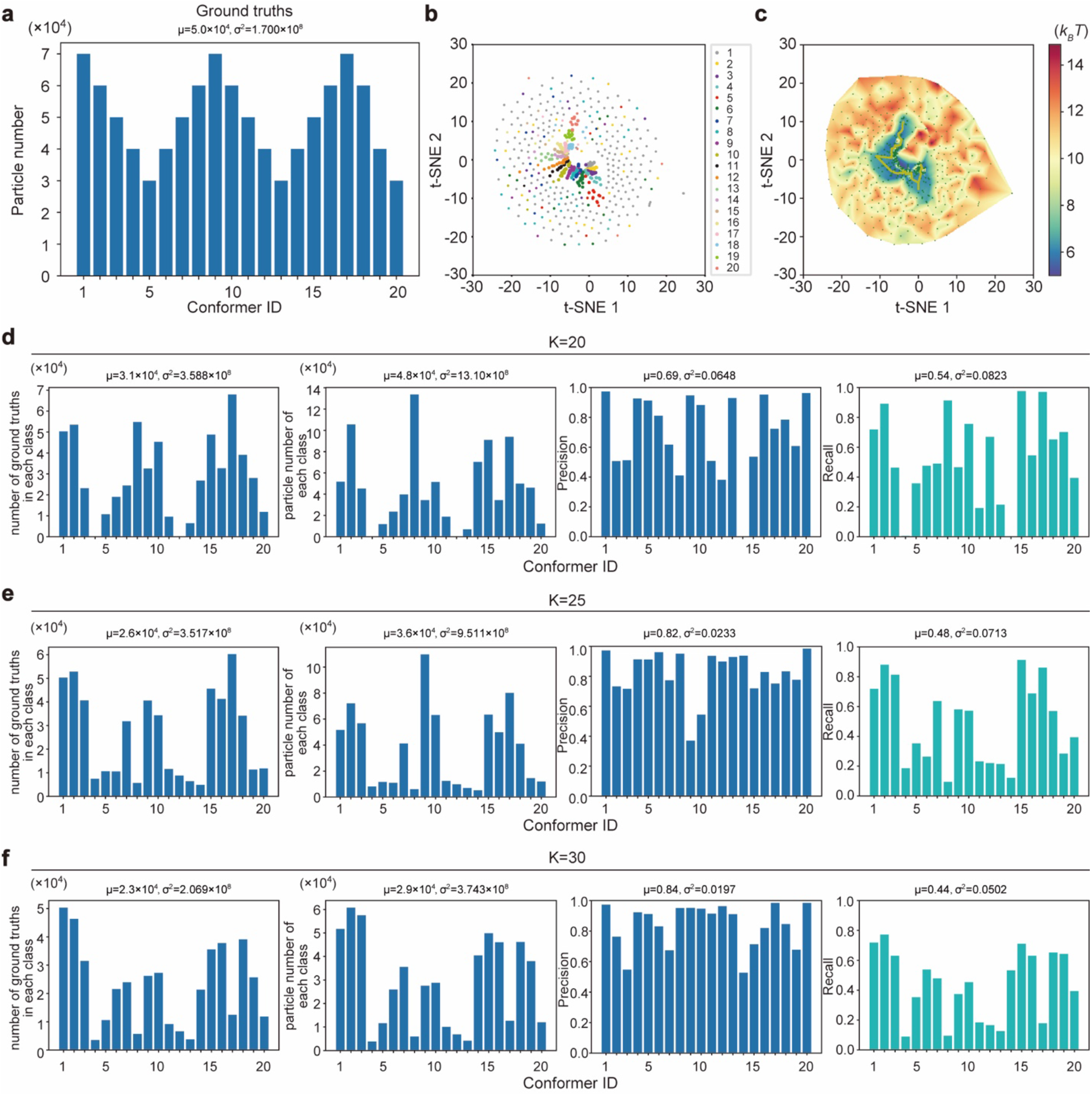
Performance evaluation of 3D classification by AlphaCryo4D on the synthetic dataset with non-uniform distribution of conformational continuum. **a**, The non-uniform distribution of 20 conformers of another simulated NLRP3 dataset at SNR of 0.01 that was used to examine the performance of AlphaCryo4D. This dataset was utilized to calculate the energy landscape shown in panels (**b, c**) and its corresponding outcome of 3D classification by AlphaCryo4D shown in panels (**d**-**f**), which tests the robustness of AlphaCryo4D in the case of non-uniform distributions of the underlying conformational states. **b**, Dimensionality reduction of bootstrapped 3D volumes and their corresponding feature maps from the simulated dataset by t-SNE. Colors of data points indicate the ground truth of their corresponding 3D volumes. **c**, Reconstruction of the energy landscape of the simulated NLRP3 dataset. **d**-**f**, The number of correctly classified particles in each class, the particle number of each class, the precision of 3D classification and the corresponding recall of AlphaCryo4D blind tests using this simulated dataset when the class number is set to 20 (**d**), 25 (**e**) and 30 (**f**).

**Supplementary Fig. 7.**
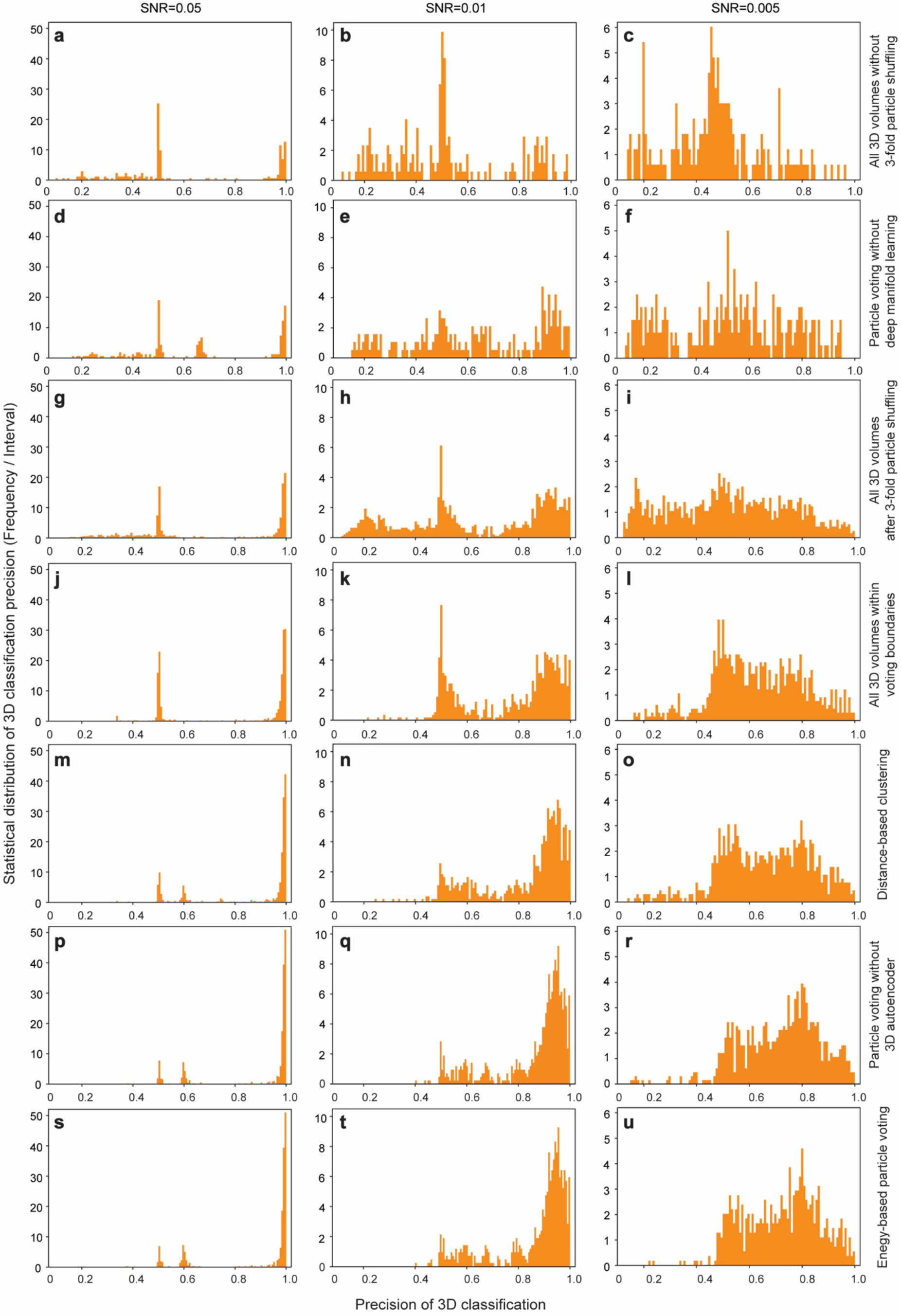
Mechanistic characterizations of the improvement of 3D classification accuracy by AlphaCryo4D using the simulated NLRP3 datasets of three typical SNRs. Left, middle and right vertical columns show the precision analysis on the simulated datasets with SNRs of 0.05, 0.01 and 0.005, respectively. In total, there are 21 conditional controls under six different algorithmic design variations being analyzed in panel (**a**)-(**r**). **a**-**c,** Statistical distribution of 3D classification precision in bootstrapped 3D volumes without particle shuffling. **d**-**f,** Statistical distribution of 3D classification precision by implementing particle voting directly on the bootstrapped 3D volumes without using deep manifold learning. The procedure of 3-fold particle shuffling and volume bootstrapping is identical to AlphaCryo4D. **g**-**i,** Distribution of 3D classification precision in bootstrapped 3D volumes after 3-fold particle shuffling in the intermediate step of AlphaCryo4D. **j**-**l,** Distribution of 3D classification precision in bootstrapped 3D volumes screened by voting boundary on the energy landscape in the intermediate step of AlphaCryo4D. **m**-**o,** Distribution of 3D classification precision after distance-based 3D clustering in the absence of energy-based particle voting with all prior steps identical to AlphaCryo4D. **p**-**r,** Distribution of 3D classification precision after particle voting by a modified AlphaCryo4D variation only without using deep residual autoencoder in the manifold embedding step. **s**-**u,** Distribution of 3D classification precision after energy-based particle voting via a complete AlphaCryo4D procedure.

**Supplementary Fig. 8.**
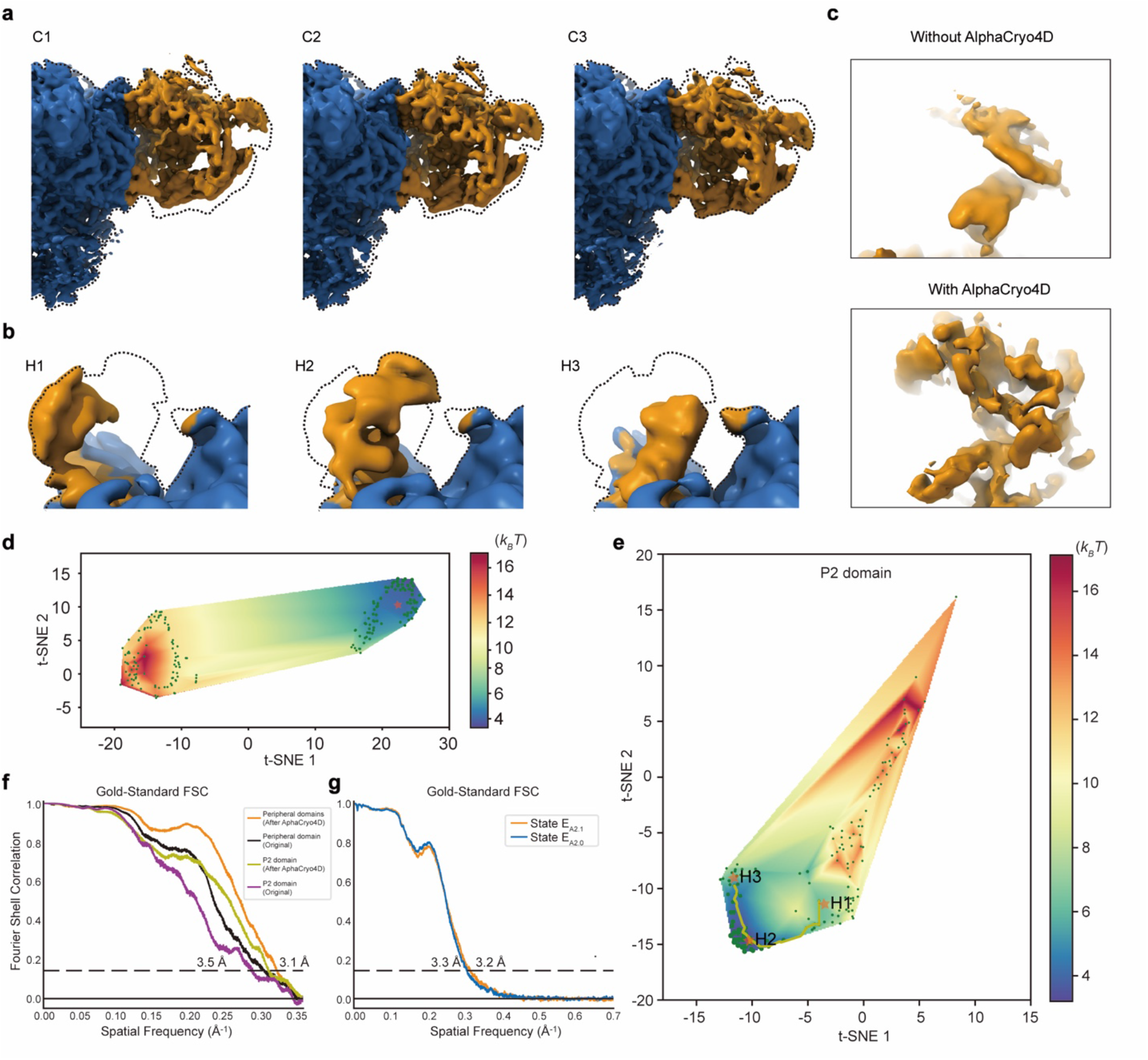
Different states of the RyR1 and resolution measure of 26S proteasome reconstructed by AlphaCryo4D. **a,** Side-by-side comparison of the cryo-EM density maps of clusters C1, C2 and C3 from the energy landscape of the peripheral domain. The dashed line indicates the outline of all density maps superimposed together. The dynamic region of the density map is colored orange, and the stable region is colored blue. **b,** Local cryo-EM density maps of clusters H1, H2 and H3 on the energy landscape of the P2 domain. **c,** Comparison of the density map of the P2 domain after AlphaCryo4D with the original state 1. **d,** Energy landscape estimated with the alignment of the peripheral domains. This energy landscape is aimed to improve the quality of the final density map. **e,** Focused energy landscape of the P2 domain of the RyR1. Clusters H1, H2 and H3 were uncovered on this energy landscape. **f,** Gold-standard FSC of the peripheral and P2 domains with and without AlphaCryo4D. **g,** Gold-standard FSC of states E_A2.0_ and E_A2.1_ of 26S proteasome reconstructed by AlphaCryo4D.

**Supplementary Fig. 9.**
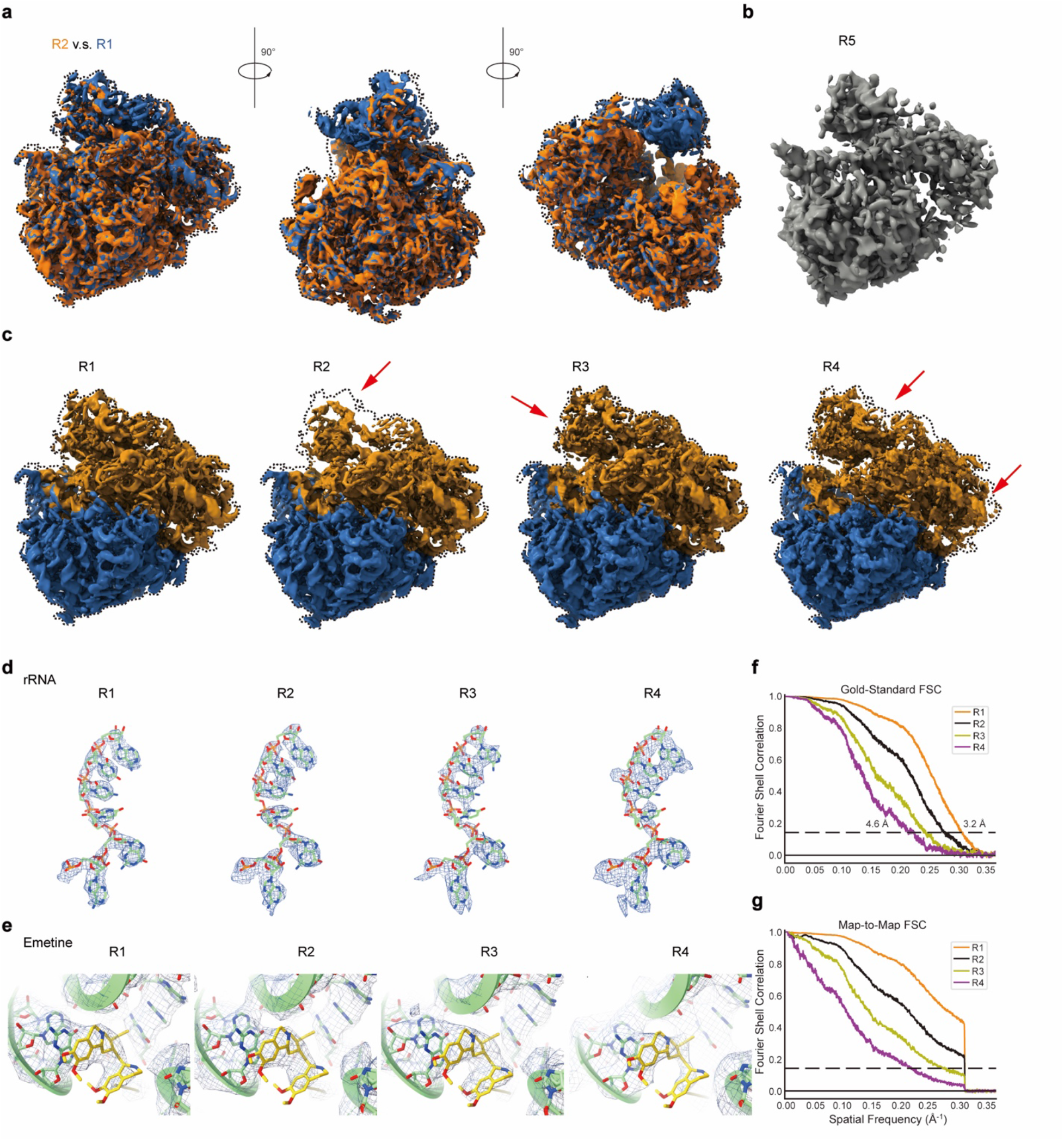
Heterogeneous analysis of the *Pf*80S ribosome reconstructed by AlphaCryo4D. **a,** Cryo-EM density map of the 40S head missing state R2 superimposed with the major state R1 from three different viewing angles. **b,** Cryo-EM density map of cluster R5 on the energy landscape (Fig. 4a). The particle number of this cluster limited the resolution of its final reconstructed density map. **c,** Comparison of the density maps of clusters R1-R4. The red arrows point to the conformational difference of states R2-R4 relative to the state R1. **d,** Comparison of local cryo-EM densities of an rRNA strand superimposed with its corresponding atomic model in four conformational states. **e,** Comparison of local cryo-EM densities of the small-molecule drug emetine bound to the ribosome, superimposed with its corresponding atomic model, in four conformational states. **f**, Gold-standard FSC of the R1-R4 states reconstructed by AlphaCryo4D. **g**, Map-to-map FSC of the R1-R4 states with the published cryo-EM structure of the *Pf*80S ribosome (EMD-2660). The cliff of the FSC curve was caused by the low-pass filtering of the previously released cryo-EM maps.

**Supplementary Fig. 10.**
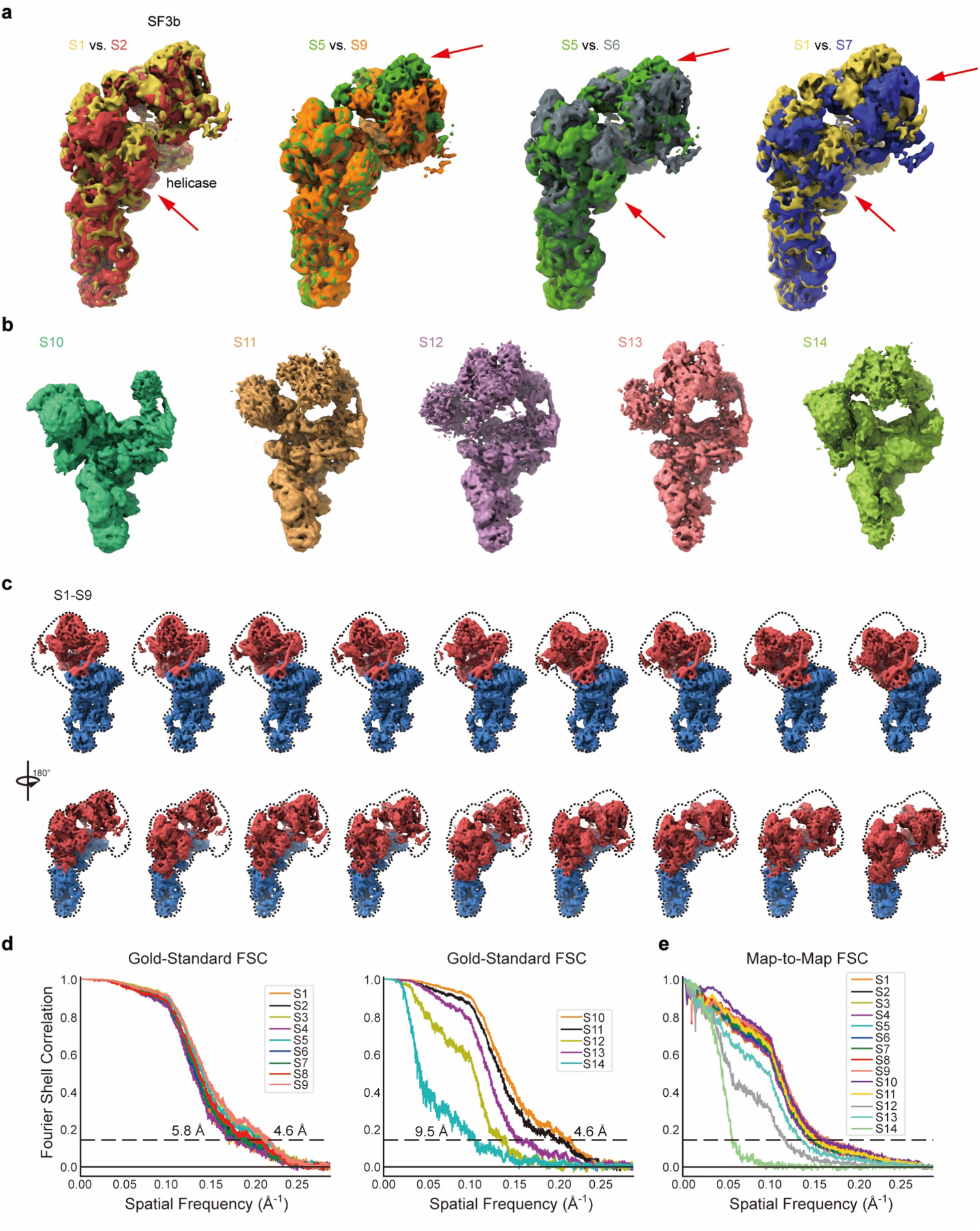
Multiple motion patterns of pre-catalytic spliceosome. **a,** Different local motion modes of the SF3b and helicase subcomplexes. The red arrows pointed the regions where the conformational changes are observed by comparing the cryo-EM densities of different states. These four comparisons exhibited the local movement of helicase (S1 vs. S2), the local movement of SF3b (S5 vs. S9), the relative motions of the SF3b and helicase subcomplexes in the opposite direction (S5 vs. S6) and the concerted motions of the SF3b and helicase subcomplexes in the same direction (S1 vs. S7), which showed that the continuous motion the SF3b or helicase subcomplexes is independent. **b,** Cryo-EM density maps of clusters S10-S14. Discrete heterogeneity is observed in these states. **c,** Cryo-EM density maps of clusters S1-S9 from other two viewing angles. The dynamic region of the density map is colored red and the rigid region is colored blue. Continuous conformational motion was observed among these states. **d,** Gold-standard FSC of the S1-S14 states classified by AlphaCryo4D. **e,** Map-to-map FSC of the S1-S14 states over the published B4 cryo-EM density map of the pre-catalytic spliceosome (EMD-3685).

**Supplementary Fig. 11.**
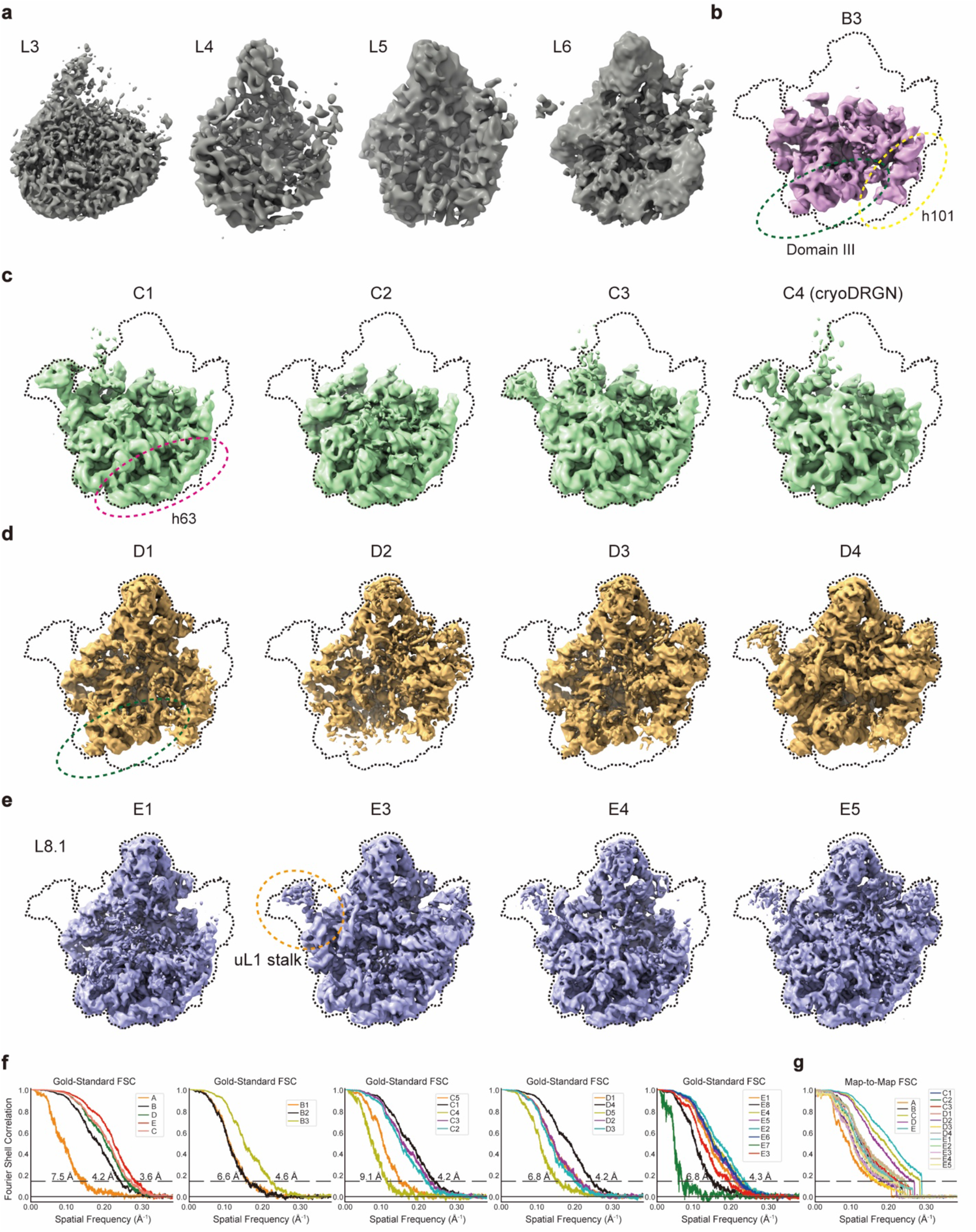
Reconstructions of bacterial ribosomal assembly intermediates by AlphaCryo4D. **a,** Cryo-EM density maps of clusters L3-L6. The classes contained particles of likely poor quality. **b,** Reconstruction of the cluster B3 from the state B energy landscape (Fig. 6d). The cluster B3 revealed the main conformation identical to state B. **c,** Reconstructions of the state C1, C2, C3 and C4 from the state C energy landscape (Fig. 6f). The state C4 contained the density of rRNA helix 68, which was previously found by cryoDRGN. **d,** Reconstructions of the state D1, D2, D3 and D4 from the state D energy landscape (Fig. 6h). **e,** Reconstructions of the state E1, E3, E4 and E5 from the state E energy landscape (Fig. 6j). **f,** Gold-standard FSC of the 5 major states (A, B, C, D and E) and 21 minor states (B1-B3, C1-C5, D1-D5 and E1-E8) reconstructed by AlphaCryo4D. **g,** Map-to-map FSC of the existing states over the published cryo-EM structures of the states A (EMD-8434), B (EMD-8440), C (EMD-8441), D (EMD-8445), E (EMD-8450), C1 (EMD-8442), C2 (EMD-8443), C3 (EMD-8444), D1 (EMD-8446), D2 (EMD-8447), D3 (EMD-8448), D4 (EMD-8449), E1 (EMD-8451), E2 (EMD-8452), E3 (EMD-8453), E4 (EMD-8455) and E5 (EMD-8456) of the bacterial ribosomal assembly intermediates.

**Supplementary Table 1.** Hyperparameters of the deep residual networks in the 3D autoencoder. Optimizer: Adam. Epochs: 50. Initial learning rate: 0.01. A factor of 0.1 and patience of 3 means that the learning rate will times 0.1 (factor) if the loss function does not improve in 3 (patience) epochs.

| Layer name | Output size<br>(simulation) | Output size<br>(RyR1) | Output size<br>(Proteasome) | Output size<br>(pf80S Ribosome) | Output size<br>(Spliceosome) | Output size<br>(Bacterial<br>Ribosomal<br>Intermediates) | Network structure |
| --- | --- | --- | --- | --- | --- | --- | --- |
| Conv1 | $200 \times 200 \times 200$ | $360 \times 360 \times 360$ | $300 \times 300 \times 300$ | $360 \times 360 \times 360$ | $320 \times 320 \times 320$ | $320 \times 320 \times 320$ | $5 \times 5 \times 5, 2$ |
| Conv2_x | $100 \times 100 \times 100$ | $180 \times 180 \times 180$ | $150 \times 150 \times 150$ | $180 \times 180 \times 180$ | $160 \times 160 \times 160$ | $160 \times 160 \times 160$ | $3 \times 3 \times 3, 2, \text{stride } 2$<br>$3 \times 3 \times 3, 2$ |
| Conv3_x | | | | | | | $3 \times 3 \times 3, 2$<br>$3 \times 3 \times 3, 2$ |
| Conv4_x | $50 \times 50 \times 50$ | $90 \times 90 \times 90$ | $75 \times 75 \times 75$ | $90 \times 90 \times 90$ | $80 \times 80 \times 80$ | $80 \times 80 \times 80$ | $3 \times 3 \times 3, 1, \text{stride } 2$<br>$3 \times 3 \times 3, 1$ |
| Conv5_x | | | | | | | $3 \times 3 \times 3, 1$<br>$3 \times 3 \times 3, 1$ |
| Conv6_x<br>(encoding) | | | | | | | $3 \times 3 \times 3, 1$<br>$3 \times 3 \times 3, 1$ |
| TransConv7_x | $100 \times 100 \times 100$ | $180 \times 180 \times 180$ | $150 \times 150 \times 150$ | $180 \times 180 \times 180$ | $160 \times 160 \times 160$ | $160 \times 160 \times 160$ | $3 \times 3 \times 3, 1, \text{stride } 2$<br>$3 \times 3 \times 3, 1$ |
| TransConv8_x | | | | | | | $3 \times 3 \times 3, 1$<br>$3 \times 3 \times 3, 1$ |
| TransConv9_x | $200 \times 200 \times 200$ | $360 \times 360 \times 360$ | $300 \times 300 \times 300$ | $360 \times 360 \times 360$ | $320 \times 320 \times 320$ | $320 \times 320 \times 320$ | $3 \times 3 \times 3, 2, \text{stride } 2$<br>$3 \times 3 \times 3, 2$ |
| TransConv10_x | | | | | | | $3 \times 3 \times 3, 2$<br>$3 \times 3 \times 3, 2$ |
| TransConv11 | | | | | | | $5 \times 5 \times 5, 1$ |

